# Spatial Deconvolution of Cell Types and Cell States at Scale Utilizing TACIT

**DOI:** 10.1101/2024.05.31.596861

**Authors:** Khoa L. A. Huynh, Katarzyna M. Tyc, Bruno F. Matuck, Quinn T. Easter, Aditya Pratapa, Nikhil V. Kumar, Paola Pérez, Rachel Kulchar, Thomas Pranzatelli, Deiziane de Souza, Theresa M. Weaver, Xufeng Qu, Luiz Alberto Valente Soares Junior, Marisa Dolhnokoff, David E. Kleiner, Stephen M. Hewitt, Luiz Fernando Ferraz da Silva, Vanderson Geraldo Rocha, Blake M. Warner, Kevin M. Byrd, Jinze Liu

**Author notes:** CORRESPONDING AUTHORS: Jinze Liu; Kevin Matthew Byrd.

## Abstract

Identifying cell types and states remains a time-consuming and error-prone challenge for spatial biology. While deep learning is increasingly used, it is difficult to generalize due to variability at the level of cells, neighborhoods, and niches in health and disease. To address this, we developed TACIT, an unsupervised algorithm for cell annotation using predefined signatures that operates without training data, using unbiased thresholding to distinguish positive cells from background, focusing on relevant markers to identify ambiguous cells in multiomic assays. Using five datasets (5,000,000-cells; 51-cell types) from three niches (brain, intestine, gland), TACIT outperformed existing unsupervised methods in accuracy and scalability. Integration of TACIT-identified cell with a novel Shiny app revealed new phenotypes in two inflammatory gland diseases. Finally, using combined spatial transcriptomics and proteomics, we discover under- and overrepresented immune cell types and states in regions of interest, suggesting multimodality is essential for translating spatial biology to clinical applications.

## INTRODUCTION

Spatial biology is a dynamic field that focuses on the precise understanding of the spatial distribution and relationship of cell types and their associated cell states within their native environments^1,2^. The field has been significantly advanced by rapidly expanding and maturing single-cell and spatial multiomics technologies, which preserves the spatial context of cellular and architectural features, deepening our understanding of cellular interactions, biological pathways, and identifying new cell types that can be used as targets to improve disease treatments and precision diagnoses^3–8^.

The current era of spatial biology, characterized by single-cell and subcellular resolution, multi-omics technologies in nature and even combined modalities on a single tissue section, demands more advanced tools for interpretation at scale^9^. Among the multi-step bioinformatics workflow to support the analysis of the multi-plex imaging data^10,11^, identifying cell types and their associated cell states remains a time-consuming and error-prone challenge due to issues related to segmentation noise and signal bleed-through, restricted sets of molecular and protein panel markers, and multimodal marker-linked datasets^12^. Traditional unsupervised clustering methods commonly used in scRNA-seq analysis operate by grouping cells based on the overall similarity of their marker profiles across the entire panel^13–17^. Their efficacy heavily relies on the presence of abundant markers capable of distinguishing cell populations, a characteristic commonly found in single cell sequencing data^18^. However, a significant challenge arises when dealing with predefined marker panels and cell types determined by as few as one marker^19^. This sparse marker set, often of only one modality, lacks power to separate expected cell population in the embedded feature space, posing a formidable obstacle for unsupervised clustering to detect all cell types especially the rare ones^20^. Even with extensive parameter tuning combined with multi-step clustering to identify cell populations of interest, the desired results remain elusive^21,22^. Deep learning algorithms are increasingly utilized in spatial ‘omics for cell type identification, but it requires comprehensive and diverse training data to improve the accuracy and applicability of deep learning models in handling the complexities of spatial multiomics^23,24^.

To address these challenges, we developed **TACIT** (**T**hreshold-based **A**ssignment of **C**ell Types from Multiplexed **I**maging Da**T**a), an unsupervised algorithm for assigning cell identities based on cell-marker expression profiles. TACIT uses a multi-step machine learning approach to group cells into populations, maximizing the enrichment of pre-defined cell type signatures from spatial transcriptomics and proteomics data (Fig. 1). Validated against expert annotation and available algorithms using five datasets from brain, intestine, and gland tissues in human and mouse, TACIT outperformed three existing unsupervised methods in accuracy and scalability. It also integrated cell types and states with a Shiny app to reveal new cellular associations in Sjögren’s Disease and Graft-versus-host Disease, highlighting its clinical relevance. Furthermore, we performed spatial transcriptomics and proteomics on the same slide, demonstrating the need for multimodal panel designs and flexible analysis pipelines to support translational and clinical research applications.

**Figure 1.**
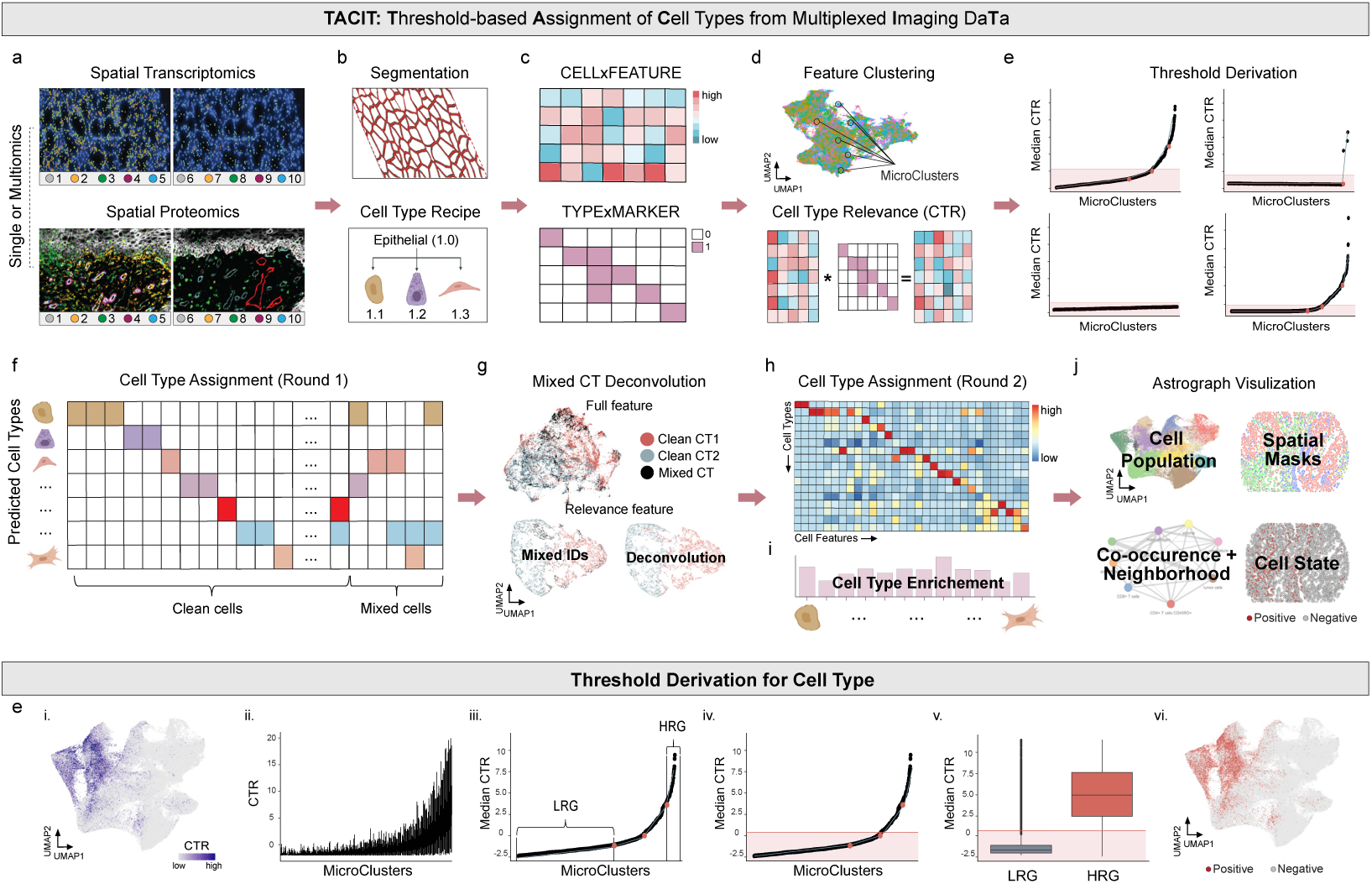
General TACIT Workflow: (a) Multiplex imaging employs both spatial proteomics (top) and spatial transcriptomics (bottom). After segmentation (b top), a CELLxFEATURE matrix is generated (c). Hierarchical cell type structures (b bottom) are formulated based on panel design, expert knowledge, and scRNA-seq marker matching, resulting in a CELLTYPExMARKER matrix (c). Cells are organized into microclusters (MCs) by a community-based Louvain algorithm, averaging 0.1%-0.5% of the population (d top). These matrices are then used to compute Cell Type Relevance (CTR) scores for all cell types across cells (d bottom). Optimal thresholds are established to classify cells as clean if they meet one threshold or mixed if multiple (e). The UMAP with all features shows no clear separation between two distinct cell types (g – top left); however, clear segregation appears when only relevant features are used in the UMAP embedding (g – top right). Mixed identities are resolved by analyzing the mode of cell types within their k-nearest neighbors (g – bottom). Validation is performed via heatmaps comparing mean marker and cell type values with the CELLTYPExMARKER matrix (h – top), and by calculating enrichment scores for each cell type (i – bottom). The UMAP plot illustrates spatial distributions with cell type annotations (j top-right) and connections of cell type clusters (j bottom-left), combining cell type and state analyses (j bottom-right). Extended details of step e: Threshold derivation extends to segmental regression on ordered median CTR scores across all MCs to identify breakpoints (i & ii), defining “low relevance group (LRG)” and “high relevance group (HRG)” (ii). The determined CTR threshold minimizes classification error, distinguishing between LRG and HRG (iv & v). Cells above the threshold are highlighted in red on the UMAP, while those below are in grey (vi).

## RESULTS

### Conceptualization of TACIT for Spatial Multimodal

To address the need for advanced spatial omics profiling, we developed an unsupervised algorithm called **TACIT** (**T**hreshold-based **A**ssignment of **C**ell Types from Multiplexed **I**maging Da**T**a). It is generally applicable to any probe-based, single-cell resolved spatial single modality or multimodal dataset (i.e., spatial transcriptomics or proteomics; Fig. 1a). Before TACIT can be employed, images containing tissues or cells are first segmented to identify cell boundaries (Fig. 1b). Features like probe intensity (protein antibodies) and count values (mRNA probes) are quantified, normalized, and stored in a single or multimodality CELLxFEATURE matrix (Fig. 1c). The TYPExMARKER matrix is derived from expert knowledge, with values between 0 and 1, indicating the relevance of markers for defining cell types (Fig. 1c).

TACIT conducts cell type annotation in two rounds. Cells are first clustered into microclusters (MCs) to capture highly homogenous cell communities with sizes averaging between 0.1–0.5% cells of the population using the Louvain algorithm (Fig 1d). In parallel, for each segmented cell, Cell Type Relevance scores (CTRs) against a predefined cell types will be calculated by the multiplication of its normalized marker intensity vector with the cell type signature vector (Fig 1d), quantitatively evaluating the congruence of cells’ molecular profile with considered cell types. The higher the CTR score, the stronger the evidence that the cell is associated with a given cell type. TACIT proceeds to learn a threshold that can separate cells into groups with strong positive signals and background noise (Fig 1e). For a specific cell type, the median CTRs across all MCs are gathered (Fig 1e^i^). The MCs are reordered by ranking its median CTRs values from lowest to highest (Fig 1e^ii^). The segmental regression model is fitted to divide the CTRs growth curve into 2 to 4 segments^25^. The two extremes of these segments represent the high relevance group and low relevance group, respectively. (Fig. 1e^iii^). A positivity threshold that minimizes the misclassification rates arising from cell outliers in both high relevance group and low relevance group is then established (Fig. 1e^iv^). Subsequently, the threshold is applied to all cells where the CTRs of cells exceeding the threshold for a specific cell type are labeled with positive, with the remaining labeled with negative (Fig 1e^v^-1e^vi^).

Cell labeling from the previous step can result in a single cell being labeled multiple cell types (Fig. 1f). To resolve the ambiguity, TACIT includes a deconvolution step (Fig. 1g) using the k-nearest neighbors (k-NN) algorithm on a feature subspace relevant to the mixed cell type category (Methods). The quality of cell type annotation is assessed by p-value and fold change, quantifying marker enrichment strength for each cell type (Fig. 1i) and visualized with a heatmap of marker expression (Fig. 1h). Following annotation, downstream analysis is performed using a custom Shiny app we generated called **Astrograph** (Fig. 1j; Methods).

### Benchmarking TACIT Against Existing Unsupervised Algorithms

We downloaded two human datasets: Colorectal Cancer (PCF-CRC; n=140-TMAs; n=235,519-cells; n=56-antibodies) and Healthy Intestine (PCF-HI; n=64-samples; n=2,603,217-cells; n=56-antibodies); both were generated using the Akoya Phenocycler-Fusion (PCF; formerly CODEX) 1.0 system for spatial proteomics^26,27^. We compared TACIT’s performance in cell type annotation against CELESTA, SCINA, and Louvain in both datasets, using original annotations as reference^13,28,29^.

In the PCF-CRC dataset, TACIT demonstrated strong consistency with reference annotations compared to existing methods. This was evident through UMAP, spatial, and heatmap visualizations of cell populations, spatial patterning, and marker expression (Fig. 2a-c). As shown in the heatmap, SCINA and Louvain missed a significant portion of rare cell types, with Louvain failing to identify 6 out of 17 types and SCINA identifying only 5 in total (Fig. 2c). TACIT achieved the highest accuracy, with weighted recall, precision, and F1 scores of 0.74, 0.79, and 0.75, respectively, significantly outperforming CELESTA, Louvain, and SCINA (p<0.05) (Fig. 2d; Extended Data 1). TACIT showed stable threshold and evaluation metrics in a bootstrap study (Extended Data 2a-d). For dominant cell types (≥1% of the population), TACIT, CELESTA, and SCINA exhibited high consistency (R=0.99) in terms cell type annotation, while Louvain slightly underperformed (R=0.95) (Fig. 2e). Both TACIT and CELESTA identified all expected rare cell types, with TACIT displaying a stronger correlation to the reference (R=0.58) compared to CELESTA (R=0.24) (Fig. 2e). Additionally, the accuracy for identifying rare cell types improved with an increasing number of resolutions (Extended Data 2e-f). Marker enrichment analysis indicated that TACIT’s annotations closely matched the signatures (Fig. 2f). Additional experiments were performed on a much larger dataset with 2.6 million cells across 40 slides, PCF-HI, derived from human intestine issues. Outperformance of TACIT over Louvain was consistently observed in overall accuracy (Figs. 2g, h, j) and enrichment strength (Fig. i), especially in its capability in identifying rare cell types (Figs. 2k). Unfortunately, both CELESTA and SCINA failed to assign a vast majority of the cells even with extensive parameter tuning (Extended Data 1).

**Figure 2:**
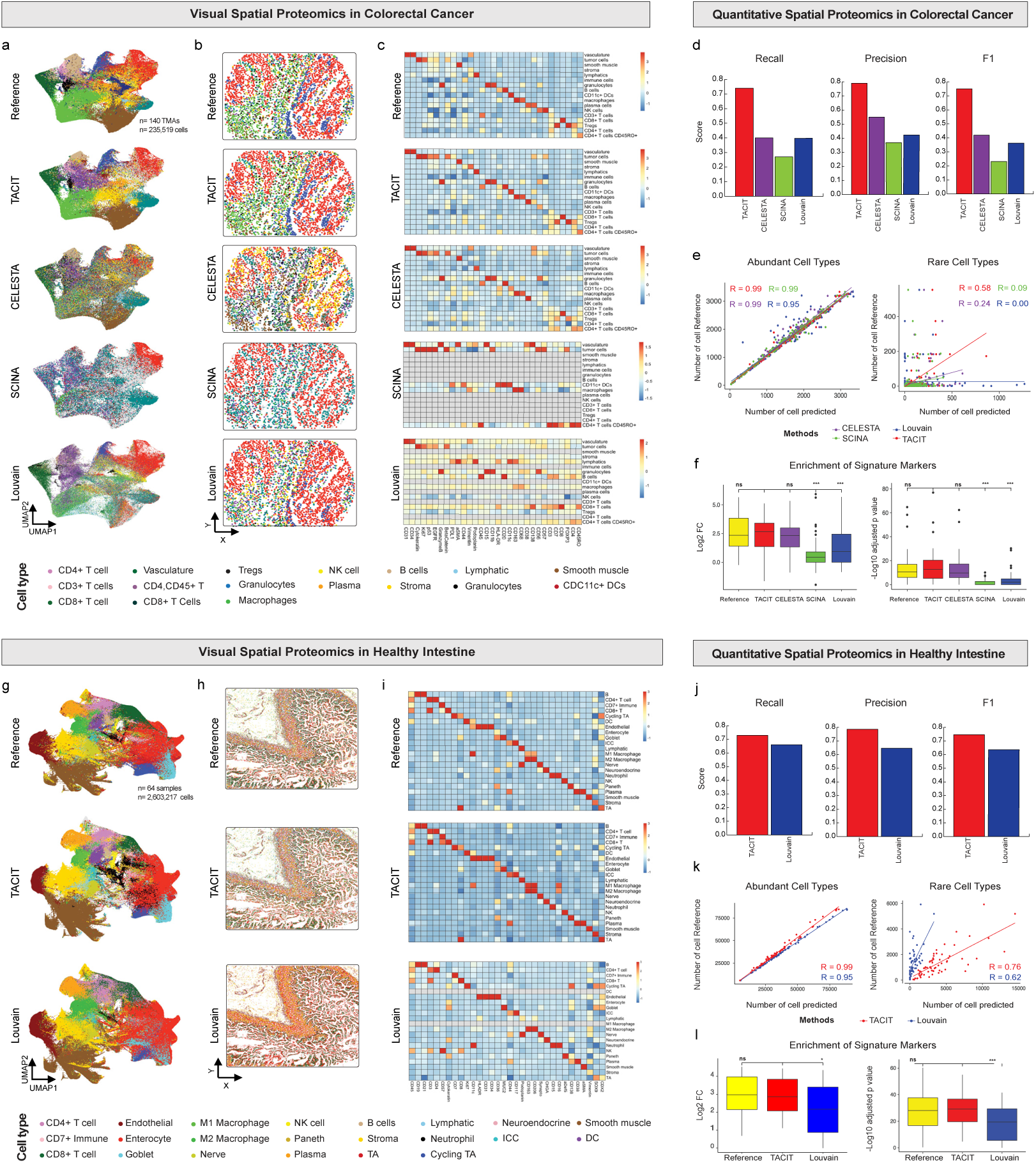
Application of TACIT on PhenoCycler data from PCF-CRC (top panel) and PCF-HI (bottom panel). (a,g) Examples of spatial plots color-coded by identified cell types, illustrating the spatial distribution and clustering of cells as determined by TACIT. These plots demonstrate how TACIT preserves the spatial structure of cell types, maintaining consistency with the reference data. (e,k) UMAP representations with cell type delineations, showing the clustering of cells in a two-dimensional space. TACIT’s UMAP plots reveal a higher degree of similarity to the reference clusters compared to other methods, indicating its superior performance in accurately identifying cell types. (f,i) Heatmaps comparing the mean marker values for each cell type identified by TACIT and other existing methods. TACIT’s heatmaps exhibit distinct and clear unique marker expressions for each cell type, with a diagonal pattern that highlights its precise cell type identification capabilities. (d,j) Recall, precision, and F1 score comparisons between TACIT (PCF-CRC: 0.74 (Recall), 0.79 (Precision), 0.75 (F1), PCF-HI: 0.73 (Recall), 0.79 (Precision), 0.75 (F1)) and existing methods, benchmarked against the reference. TACIT consistently outperforms other methods, achieving higher recall, precision, and F1 scores, which underscores its accuracy and reliability in cell type identification. (e,k) Correlation plots illustrating the relationships between different cell type identification methods for both abundant cell types and rare cell types. TACIT shows strong correlations with the reference data, particularly for rare cell types (PCF-CRC: R=0.58, PCF-HI: R=0.76), where it demonstrates a higher degree of similarity in cell type identification compared to other methods. (f,l) Intensity comparison of unique markers between TACIT and existing methods. TACIT displays significantly different enrichment scores, particularly when compared to methods like Louvain (PCF-CRC & PCF-HI: p-value<0.05) or SCINA (PCF-CRC: p-value<0.05), indicating its enhanced ability to identify and distinguish unique cell markers.

To evaluate TACIT’s performance on spatial transcriptomics data, we applied it to a published MERFISH dataset from the murine hypothalamic preoptic region of the brain (n=36-samples; n=1,027,848-cells; n=170-ISH panel)^30^. TACIT achieved significantly higher weighted recall (0.85), precision (0.87), and F1 scores (0.87) than Louvain (Extended Data 3a). Both methods showed high correlation with the reference for dominant cell types (R=0.99), but TACIT achieved higher correlation for rare cell types (R=0.94) compared to Louvain (R=0.64; Extended Data 3b). Spatial and UMAP plot demonstrated that TACIT’s cell type identification closely matched the reference, with stronger and more distinct expression signatures than Louvain (Extended Data 3c-f). These results highlight TACIT’s effectiveness for spatial transcriptomics, providing reliable cell type identification for both abundant and rare populations.

### Applying TACIT to unpublished single modality spatial transcriptomics with linked scRNAseq

Next, TACIT was applied to an unpublished Xenium dataset (PI: Warner, NIH/NIDCR; n=21-patients; n=∼360,000-cells; n=280-ISH panel) across 24 cell types. We compared TACIT against two annotation approaches: Seurat with label transfer from scRNA-seq data (Seurat transfer), and Louvain^29,31^. Signature lists for TACIT were created from the top five most enriched genes in each annotated cluster in the Seurat transfer result^32^. While the UMAP plot shows overall consistency in cell type annotation across the three methods, TACIT’s annotation excels in clear distinctions among three subtypes of acinar cells (Fig. 3a), corroborated by biologically meaningful spatial arrangement of these subtypes (Fig. 3b). TACIT demonstrated higher enrichment of signatures than both Louvain and Seurat transfer, with all cell types identified (Fig. 3c, h, I, g). Zooming into specific subtypes, TACIT clearly distinguishes ductal progenitors and ductal cells, while Seurat transfer labeled them all as "ductal cells" and Louvain showed mixed annotations (Figs 3d). TACIT also identified four subsets of T cells (CD4+, CD8+, CD8+ Exhausted, and Progenitors), which Louvain missed (Fig. 3e). This is a critical population to identify for a disease like autoimmune diseases like Sjögren’s because T progenitors are crucial for maintaining immune tolerance, making them vital targets for therapeutic strategies and clinical applications in the future^33^. Overall, TACIT showed a strong correlation with scRNA-seq (R=0.84), higher than Seurat transfer (R=0.49) and Louvain (R=0.69) (Fig. 3f).

**Figure 3:**
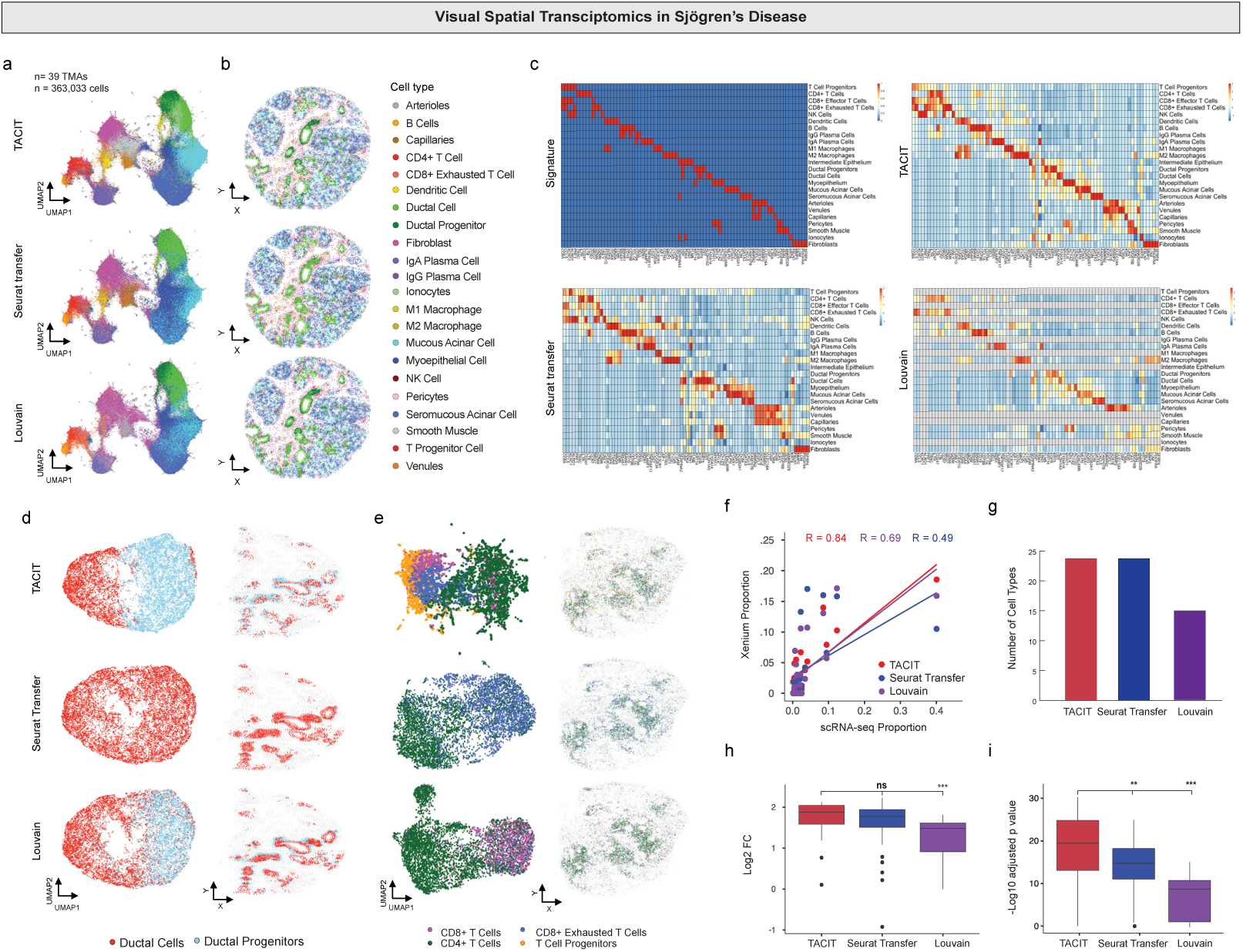
Application of TACIT on Xenium data. (a) UMAP and (b) spatial plots color-coded by identified cell types. The UMAP plots demonstrate TACIT’s ability to cluster cells accurately, showing a clear separation of different cell types. Notably, epithelial such as mucous acinar, myoepithelial, and seromucous acinar cells form more distinct and clear clusters under TACIT’s annotation compared to Louvain and Seurat Transfer methods. The spatial plots further illustrate the spatial distribution of these cell types, maintaining the structural integrity and spatial organization consistent with the reference data. (c) Heatmaps depicting cell types and markers between TACIT, Louvain, Seurat transfer, and the signature matrix. TACIT’s heatmaps present clear and distinct patterns, highlighting its precise identification of cell types and markers. This clarity is especially notable when compared to the other methods, which show less distinct marker expressions. (d-e) UMAP plots with low granularity cell types across the three methods. TACIT’s enhanced capabilities are further exemplified by its identification of rare and diverse cell types, such as duct cells and duct progenitors, as well as various T cell types including CD4, CD8, CD8 exhausted, and T cell progenitors. (f) Correlation plot of cell type proportions between the three methods in Xenium, compared with scRNA cell type proportions. TACIT shows a higher correlation (Spearman Correlation, R=0.84) with scRNA cell type proportions, indicating a more consistent and reliable identification of cell types. In contrast, Seurat transfer and Louvain show lower correlations of 0.49 and 0.69, respectively. (g) TACIT and Seurat transfer able to find all the cell type matches with scRNA. (h-i) Intensity comparison of unique markers between TACIT and existing methods. TACIT exhibits a higher intensity of unique marker expressions compared to Louvain, with a log2 fold change (p-value<0.05), and shows significant performance over Louvain and Seurat transfer, with a -log10 adjusted p-value (p-value<0.05).

### Applying TACIT to unpublished same-slide spatial proteomics and transcriptomics

To achieve detailed cell type annotation in spatial multiomics, we linked spatial proteomics (PI: Byrd, ADA Science & Research Institute; PCF 2.0; 36-antibody panel; Fig. 4a) and transcriptomics (Xenium; 280-ISH panel; Fig. 4b) on the same slide using segmentation mask transfer. This captured single-cell data for both TACIT and Louvain (see Methods; n=6-samples; 424,638-cells). Cellenics (now Trailmaker) was used to generate cell type signatures (Extended Data 4). Applied to minor salivary glands affected by Graft-versus-Host Disease (GvHD), TACIT identified significantly more cell types than Louvain in both datasets (Extended Data 5,6; Figs. 4c-e). Louvain missed key cell types like vascular endothelial cells and Tregs. The reconstructed slide showed high immune cell density in the periductal region, indicating GvHD-associated immune infiltration (Fig. 4d). Compared to the pathologist’s annotations, TACIT had a lower error rate than Louvain across all cell types (Figs. 4f).

**Figure 4:**
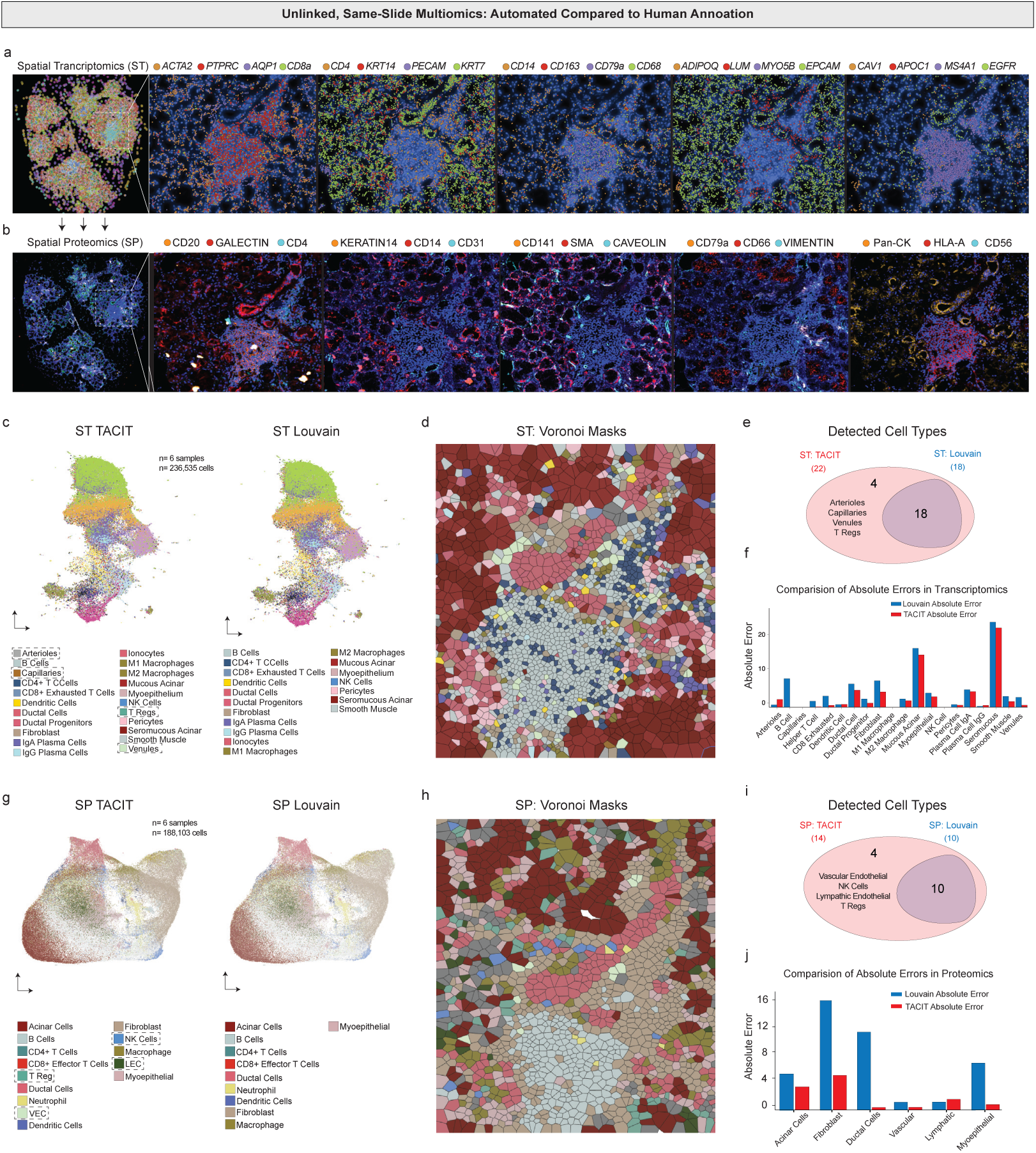
Single-Slide Spatial Multiomics Annotation using TACIT. (a) Spatial Transcriptomics – A Xenium experiment was conducted on minor salivary glands of GVHD patients using a 280-gene panel focusing on structural and immune cells. The inset shows a high-density immune area and the overlay of representative structural, immune, and cell state transcripts in the area of interest. (b) Spatial Proteomics – A post-Xenium Phenocycler Fusion experiment was performed on the same slide, using a 36-antibody panel targeting structural and immune cells. The segmentation mask was shared between both experiments to extract spatial single-cell data. (c) The UMAP of the Xenium data using TACIT and Louvain shows a higher granularity in the annotations made by TACIT. The cell types identified solely by TACIT are highlighted in the cell type annotation (*arrows*). (d) Voronoi plot showing TACIT’s annotation reconstruction of a GVHD case. The inset reveals the heterogeneity of cells detected in a high-density immune infiltrate. (e) Venn diagram showing the matched and unique cell types detected by each tool in the spatial transcriptomics experiment. TACIT identified 22 cell types, with 4 not matched by Louvain. All cell types detected by Louvain were also detected by TACIT. (f). The absolute error of cell assigns compared with human pathologist evaluation, for each cell type using TACIT and Louvain. (g) The UMAP of the Phenocycler Fusion data using TACIT and Louvain shows a higher granularity in the annotations made by TACIT. The cell types identified solely by TACIT are highlighted in the cell type annotation (*arrows*). (h) Voronoi plot showing TACIT’s annotation reconstruction based on a spatial proteomics assay of a GVHD case. The inset shows the heterogeneity of cells detected in a high-density immune infiltrate at a lower resolution compared to the spatial transcriptomics. (i) Venn diagram showing that TACIT recognized and assigned 18 cell types, with two structural and two immune cell types uniquely detected by TACIT. (j) The absolute error of cell quantity signatures using a spatial transcriptomics assay, compared with human pathologist, for each cell type using TACIT and Louvain.

In spatial proteomics, TACIT again identified more cell types than Louvain (Figs. 4g,i), matching the spatial transcriptomic assignments and confirming GVHD-associated immune infiltration (Fig. 4h). TACIT uniquely identified vascular and lymphatic endothelial cells, Tregs, and NK cells (Fig. 4i). TACIT also had a lower mean error in annotating structural cell types, while Louvain over-assigned prevalent types like fibroblasts and ducts (Fig. 4j). Vascular and innate cell types are crucial markers for understanding salivary gland parenchymal changes in GVHD; in particular, NK cells can contribute to the severity of GVHD by directly killing host cells and releasing inflammatory cytokines such as IFN-γ and TNF-α^34^. This highlights the importance of selecting the right tool for accurate cell annotation, from basic to clinical studies involving human subjects.

### Testing TACIT in linked spatial proteomics and transcriptomics ROIs

Because specific ROIs are often used for diagnosis or understanding disease pathophysiology, we decided to evaluate TACIT’s performance in confined areas. We selected nascent tertiary lymphoid structures (TLS) from GVHD for this application. TLSs pose unique challenges for spatial biology due to potential segmentation issues as they are highly concentrated with immune cells with large nuclei and little cytoplasm around diverse structural niches (epithelial, fibroblast, and vasculature)^35^. We applied a segmentation pipeline using a human-in-the-loop Cellpose3 model and still found areas in the TLS in both proteomic and transcriptomic space where signals like those for B Cells (protein: CD20; mRNA: *MS4A1*) are misappropriated after segmentation Fig. 5a)^36^.

**Figure 5:**
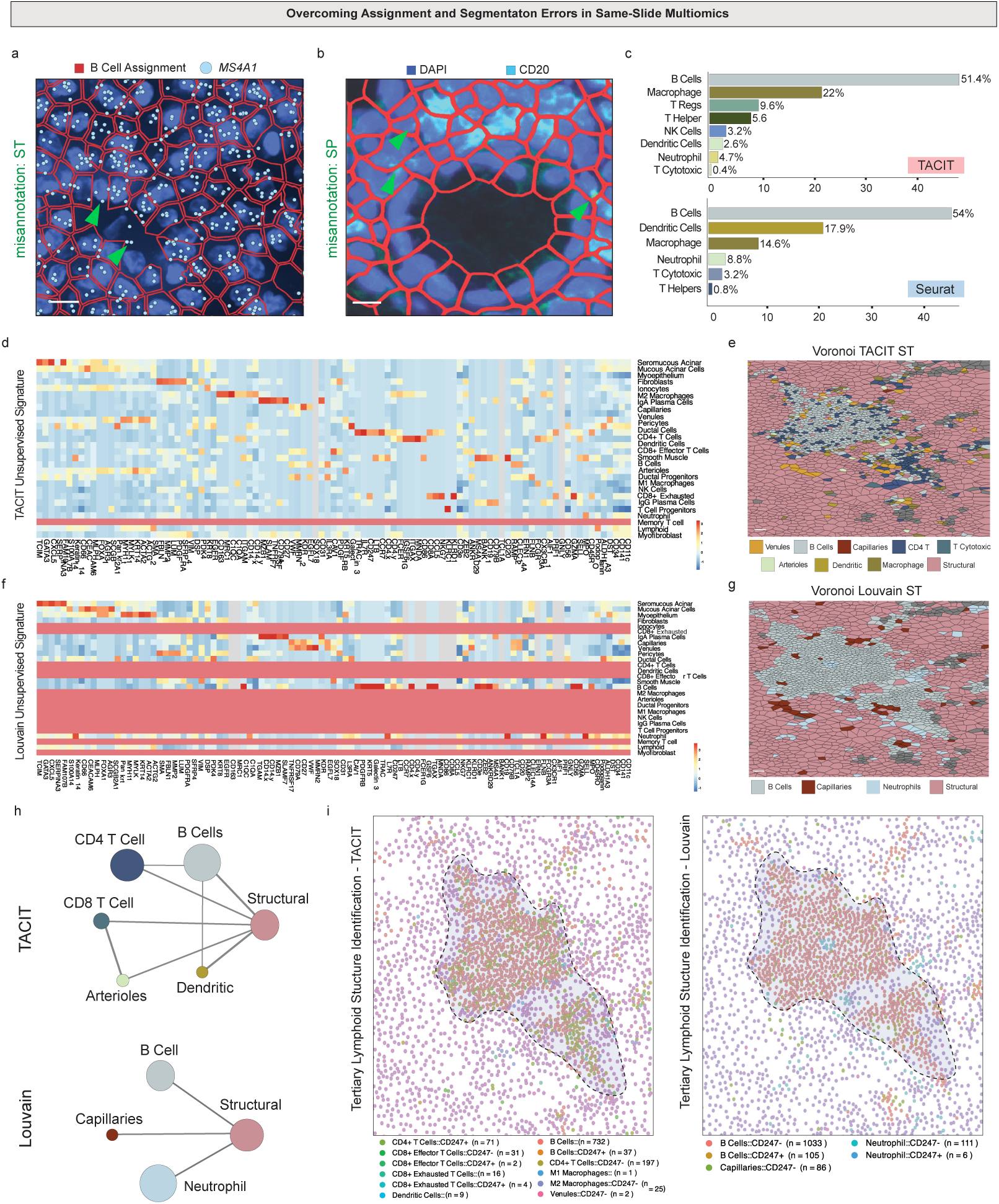
Application of TACIT in a Multimodal Single-Slide Tertiary Lymphoid Structure. (a) Spatial transcriptomics and proteomics assays were used for segmentation to extract spatial single-cell data. The segmentation mask was transferred from experiment to the another, even then it can present bleed-through of markers between cells; proteomics data can show immunofluorescence markers staining the edges of adjacent B cells (arrows). The same issue can occur with transcript probes being detected outside the cell boundary, as shown in a tertiary lymphoid structure (TLS) in a GVHD minor salivary gland where the *MS4A1* gene is detected outside of B cells. (b) TACIT and Louvain have different performances when analyzing high-density immune areas of interest, such as a TLS. The immune cell proportion identified by TACIT showed a more detailed population of cells expected in a TLS compared to Louvain. (c) A heatmap shows the genes and proteins used to create cell signatures by TACIT and Louvain. Using the same list of genes, TACIT outperforms Louvain in terms of clear markers for each cell type and cell recognition in a high-density immune cell area. (d) Voronoi plots illustrate how different assignments can create varying outcomes and analyses. The TACIT reconstruction using the signature list shows a heterogeneity of immune cells surrounded by small vessels and antigen-presenting cells, as expected in a TLS. Louvain presented a lower resolution of cell recognition, combining all immune cell types into just one innate and one adaptive immune cell type. (e) Downstream analysis can be impacted by using different tools to assign cells in multi-omics spatial assays. The neighborhood analysis presented by a Delaunay triangulation shows the expected proximity of cells in a TLS, such as B cells and dendritic cells with small vessels and T cells using TACIT. Louvain presented unilateral interactions, all related to the structural cell types, which were the most abundant cell type in the ROI analyzed. (f) The use of a single-slide spatial proteomics and transcriptomics opens the possibility of finding cell types and assigning chemokines, interleukins, and immune checkpoints to each cell type. This not only detects cellular patterns but also begins to explore spatial cell-cell communication validation and interactions. The ROI reconstruction using TACIT showed *CD247* assigned to T cells, B cells, and macrophages, whereas Clustering’s signature was unique to B cells and surrounded by capillaries with no other interactions.

TACIT’s ability to deconvolve mixed cell phenotypes helps overcome segmentation errors. Within the TLS, TACIT identified more adaptive and innate immune cell types than Louvain, including Regulatory T Cells and NK Cells (Fig. 5b). Louvain detected fewer cell types with less distinct markers per cell type (Fig. 5c). In Voronoi reconstruction, Louvain identified TLS mainly composed of B Cells, while TACIT showed primarily T cells surrounded by small vessels (Fig. 5d). Neighborhood analyses using Delaunay Triangulation and receptor-ligand pairs revealed different TLS phenotypes. TACIT showed expected relationships, such as proximity between dendritic cells and T cells, while Louvain showed structural-to-structural cell relationships (Fig. 5e). TACIT identified key markers for T cell exhaustion (PD-1/PD-L1 interactions) and small vessels essential for immune cell recruitment, while Louvain failed to detect vascular cells and showed less granularity in receptor-ligand assignments. This analysis demonstrates that niche- and disease-level phenotyping can be effectively captured using TACIT’s workflow.

### Multimodal Cell Identification with TACIT

After collecting spatial transcriptomics (Xenium) and spatial proteomics (PCF) data, we used the same segmentation masks from Xenium on the PCF data, ensuring matched cell IDs for direct comparisons (see: Methods and Fig. 6a). This alignment allowed us to create a cell-by-protein and gene matrix for each cell, capturing both antibody intensities from PCF and count values from Xenium (Figs. 6b,c). Using TACIT, which incorporates marker signatures from both PCF and Xenium, we accurately identified cell types; other algorithms could not handle the multimodality for these assays. For the first time, the correlation of marker intensities between PCF and Xenium for immune cell markers was significantly lower than for structural cell types (p<0.0001) (Fig. 6d). Consequently, using the full marker panel on ROIs with many immune cells, the agreement between cell type identifications using only PCF markers versus only Xenium markers was about 34% (Fig. 6e and Extended Data 6a). However, focusing on markers common to both PCF and Xenium increased the agreement to 81% (Fig. 6f and Extended Data 7a). The proportion of cell types was high in the TLS between Xenium and PCF with higher agreement when using common markers (Figs. 6g,h). Importantly, for structural cell types like vascular endothelial cells (VEC) using our panel, they remained challenging to identify (see Fig. 4).

**Figure 6.**
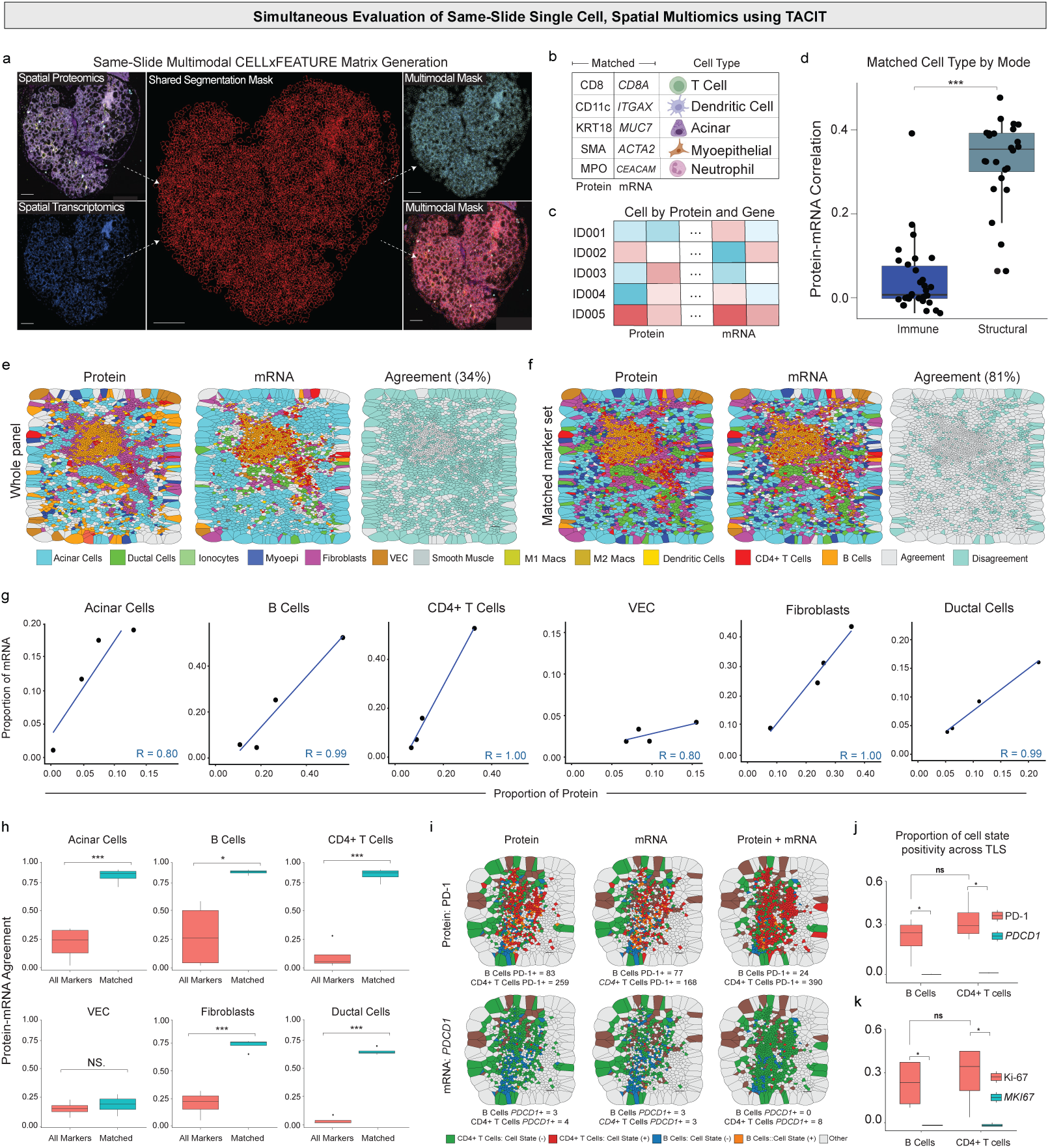
Multimodal analysis using ST and SP in a single slide. (a) Two assays were combined on the same slide and section: Phenocycler Fusion (SP) and Xenium (ST). These were performed using a 36-antibody panel and a 280-gene panel. A segmentation mask was created using a human-in-the-loop approach and inputted into the Xenium Ranger. This mask was then transferred to the SP assay, maintaining cell IDs between the two experiments.(b) After segmentation, a matrix was extracted containing the pixel values of each immunofluorescent channel from the SP and the transcripts per cell from the ST. (c) This cell-by-feature matrix was then normalized and cell-assigned using TACIT. (d). The matched number of cells assigned by the SP and ST assays was quantified to evaluate the correlation in cell assignment for each major cell type – structural and immune cells. The correlation for structural cells using all transcripts and proteins was 0.37, and for immune cells, it was 0.01. (e). After the initial annotation, specific cell markers were used to assign cell types that had both protein and transcript designations in the proteomics and transcriptomics assays. The masks of cells annotated in three different ROIs with a high density of immune cells showed 34% agreement when using all markers. (f). A smaller subset of matched protein and RNA panels was utilized to improve agreement. The Voronoi mask showed better convergence in cell type annotation, increasing cell ID matching to 81%. (g-h) The difference in annotation by each approach for each of the six cell types selected using matched protein and RNA markers showed an improvement in cell assignment, with the proportion of the cell types. (i). After multimodal cell assignment, TACIT was also able to provide cell state markers for each cell. PD-1 and *PDCD1* were used to understand the ratio of transcripts and proteins in high-density immune cell ROIs. The presence of these two markers was analyzed using SP alone, ST alone, and the two assays combined. (j) The proportion of positivity cell state in mRNA such as *PDCD1* and *MKI67* are significantly lower than PD-1 (p-value<0.05) and Ki67 (p-value<0.05) in protein for B cells and CD4+ T cells across TLS.

For effective clinical translation, it is crucial to accurately assign both spatial cell identity and state. To address this, we tested PDCD1/PD1, a key component of the immune checkpoint inhibitor (ICI) pathway. The differences observed across all three recipes—unimodal and multimodal—highlight the importance of understanding which factors are truly critical for patient outcomes, especially as they vary with spatial scales in cell number and sample number. Comparing the same markers across both technologies revealed differences in cell states, particularly between PD-1 and *PDCD1* across all four TLS (Fig. 6i and Extended Data 7b). These results were statistically significant for B cells and CD4+ T Cells (Fig 6.j). The same trend followed for cell cycling marker Ki-67/*MKI67* (Fig. 6k). This is clinically relevant because accurately predicting the cell cycle and PD-1 expression in B cells and CD4+ T cells is crucial for optimizing immunotherapy, as it helps identify which patients will benefit most from treatments like checkpoint inhibitors.

## DISCUSSION

Identifying cell types in multiplex imaging-based spatial omics data remains challenging with current technologies. In contrast to unsupervised clustering algorithm requiring extensive manual curation, TACIT automates cell type annotation, emulating manual gating with scalability and precision. TACIT achieves detailed phenotyping based on the multiplex panel design and excels in dominant and rare cell populations without bias. The success of TACIT can be attributed to the usage of cell-type specific features, initially evaluating cell type-specific markers, then performing mixed cell deconvolution within only relevent subspace, crucial for identifying cell types in spatial transcriptomics and proteomics platforms where specific features are sparse.

Our benchmarking of TACIT on three public spatial omics datasets totaling nearly 4.6 million cells across 51 cell types demonstrating its broader applicability as an assay-, species-, organ- and disease-agnostic tool for cell type annotation. Our application of TACIT to the Xenium dataset initially annotated by scRNA-seq data through label transfer further demonstrated TACIT’s effectiveness in refining cell type annotations following the discovery of cell type specific markers through existing exploratory analysis.

The combined analysis of spatial multiomics datasets in GVHD revealed the importance of integrating spatial transcriptomics and proteomics for deep phenotyping. PCF and Xenium data differ in that PCF provides continuous values while Xenium provides count data, and there is often a lack of correlation between corresponding markers, especially structural ones^37^. Despite these challenges, TACIT supports both data types, enabling high-quality targeted deep phenotyping and comparative analysis. This facilitates the combination of datasets to uncover important cellular neighborhoods and characterize cell states across modalities.

Proper application of TACIT requires sufficient sampling of cells with abundant background signals to derive relevant thresholds, making it less effective when focusing on small regions with few cells. Additionally, TACIT may leave some cells unassigned due to poor marker intensities, inaccurate segmentation, or the presence of novel cell types. Further investigation of unannotated cells to support novel cell type discovery can be achieved by a variation of TACIT capable of identify cell groups exhibiting combinations of positive markers.

By providing detailed cell type annotations and uncovering rare cell populations, tools like TACIT enables the identification of unique cellular neighborhoods and their interactions, which is critical for understanding disease progression and therapeutic response in the near future as part of clinical research and ultimately, precision clinical care. As TACIT continues to evolve, its application in personalized medicine could lead to the development of tailored treatment regimens based on the specific cellular composition and state of individual patients’ tissues, improving outcomes and reducing adverse effects.

## METHODS

### CELLxFEATURE matrix

Let *M* be a set of markers used in a spatial omics panel, |M|=m, and *N* be the set of cells of size *n* captured in a tissue slide. Let *A*_*n*×*m*_be the CELL by FEATURE information captured in the spatial omics experiment following cell segmentation process. For spatial proteomics such as PhenoCycler, entry *a*_*ij*_ in the matrix A represents the z-normalized intensity value indicating the Intensity level of a specific marker *j* within cell *i*. In the context of spatial transcriptomics such as Xenium or MERFISH/MERSCOPE, *a*_*ij*_ reflects the log-normalized of the count of transcripts for each gene.

### Cell type signature matrix

Let Τ be a set of cell types, |Τ| = t, to be captured by the panel. We define a cell signature matrix *S*_*m*×*t*_ of markers that define individual cell types, where each element *s*_*i*j_ in *S*

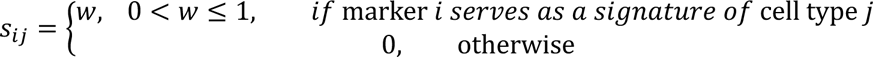

The value *w* indicates the importance of a specific marker in defining a cell type. If such information is not available, *w* is set to 1 by default.

### Cell type relevance matrix

Let Γ denote a cell type relevance matrix, with dimension *n* × *p*, where *n* is the number of profiled cells, and *p* is the number of cell types included in the panel. The cell type relevance (CTR) score is computed using the formula:

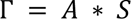

where each element in Γ provides a quantitative measure of a cell’s relevance to a specific cell type. By summing up the relevant markers’ intensity values weighted by their importance (set to 1 by default), we can directly measure a cell’s marker intensity profile alignment with the expected cell type signature. For each cell type, a cell with higher CTR score suggests a stronger association between the observed marker intensities with the expected signature of a specific cell type, indicating a higher likelihood that the cell belongs to that cell type.

### Micro-clustering

Louvain clustering method from the Seurat version 5 toolkit is applied on the CELLxFEATURE matrix *A* to conduct the fine-grained clustering of cells^31^. The resolution of the clustering is set high enough so that the average number of cells per cluster remains between 0.1% to 0.5% cells of the entire population. We refer to the resulting clusters as a collection of microclusters (MCs) denoted as Φ = {*c*_1_, *c*_2_, …, *c*_!*κ*_}. These microclusters are expected to be highly homogeneous, capturing a group of cells with highly similar marker profiles and thus with high likelihood to represent cells of the same cell type. The distribution of marker values across all markers in Φ will be used to approximate the variations of marker values across the diverse cell populations they represent.

### Segmented regression model

Next, to identify MCs with distinct cell type relevance, we employed segmented regression model aiming to identify specific breakpoints at which the relationship between the MCs changes^25^. For any given cell type, the median CTR scores across all *k* MCs are calculated and stored as a vector *z* = (*z*_1_, *z*_2_, … *z*_*κ*_) = (*r*_1_, *r*_2_, …, *r*_*κ*_) be a vector where *r*_*i*_ is the rank of *z*_*i*_in *z*. Next, a segmental regression model is fitted with *z* being the dependent variable and *r* as the predictor to identify breakpoints that divide the data into distinct linear segments.

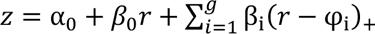

Where:

- *α*_0_ represents the intercept of the linear model,
- *β*_0_ represents the slope of the linear segment before the first breakpoint,
- *β*_*i*_ represents changes in slope at the breakpoint *i*,
- *g* represents number of breakpoints,
- φ_i_ represents the optimal location of breakpoint *i*,
- (*r* − φ_i_)_+_ defined as max(0, *r* − φ_i_) for breakpoint *i*.

Our proposed method aims to obtain an optimal fitting by allowing a maximum three breakpoints. This is determined by the minimal Akaike Information Criterion (AIC) score achieved among the three models the three models (g=1, 2 and 3)^38^. The breakpoints from the optimal model are then utilized to categorize clusters into either "low" or "high" relevance groups, Φ_L_ and Φ_H_, respectively. Specifically, the MCs ranking below the lowest breakpoint are classified as Φ_L_ = {*i* |*r*_*i*_ ≤ φ^1^, 1 ≤ *i* ≤ *κ*}, where *r* is the vector containing the rank positions of MCs. Correspondingly, the MCs ranking above the highest breakpoint are considered as high relevance group Φ_H_= {*i* | *r*_*i*_ ≥ φ^max(g)^, 1 ≤ *i* ≤ *κ*}.

### Optimal threshold

Next, an optimal CTR threshold to differentiate positive and negative cells of a given cell type is determined as follows. Let *C*_*L*_ denote the set of cells that belong to MCs within Φ_L_, formally defined as *C*_*L*_ = ⋃_*i* ∈ ΦL_ *c*_*i*,_ *c*_*i*_ ∈ Φ. Similarly, *C*_*H*_ is the set of cells that belong to MCs within Φ_H_, defined as *C*_*H*_ = ⋃_*i* ∈ ΦH_ *c*_*i*_, *c*_*i*_ ∈ Φ. Each MC encompasses a range of CTR scores, suggesting that even within a highly homogeneous cluster, there is relatively broad range of marker intensity. The preferred threshold minimizes the misclassification rate between the two relevance groups. This optimization problem aims to find a threshold (*θ*) that minimizes the number of cells in the low relevance group *C*_*L*_ with CTR scores exceeding the threshold, and the number of cells in the high relevance group *C*_*H*_ with CTR scores lower than the threshold. The grid search with this objective function can be expressed with the formula:

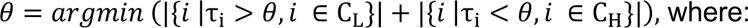

- θ represents a desired optimal threshold for a given cell type,
- τ_i_ is the CTR score for cell *i*,

### Cell Type Categorization

After determining an optimal threshold of CTR score for each cell type, cells exceeding this threshold are marked as positive, while the rest are marked as negative. Applying this threshold to each cell type results in a binary matrix Β of dimension *n* × *p*, with 1 indicating a cell is positive or 0 indicating negative. Based on the positivity of individual cells across cell types, cells are categorized into three distinct sets:

1. Clean cells: The set of cells classified as positive for exactly one cell type.
2. Mixed cells: The set of cells classified as positive in more than one cell type, suggesting a blend of characteristics from multiple cell types.
3. Unknown cells: The set of cells that are not classified as positive for any cell type.

### Deconvolution of Mixed Cells

The set of mixed cells undergoes a process of cell type deconvolution to assign each cell to its final cell type. This step leverages two outcomes from the previous step. Firstly, a significant portion of cells classified as clean cells in each individual cell type may now serve as anchor cells to resolve the cells with mixed identities. Secondly, even though more than one identity is assigned as candidates for mixed cells, a vast majority of cell types are recognized as irrelevant and will be eliminated from further consideration. So are the markers from irrelevant cell types, allowing the classification algorithm to focus on the relevant markers to resolve the confusion while avoiding distractions from irrelevant markers.

Let *ξ*, *ξ* ⊂ Τ, be a combination of cell types deemed positive in a set of cells, denotes as 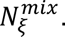 Additionally, all the clean cells positive in each of cell types in *ξ* are also extracted, denoted as 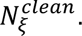 Let *M*_<_ be the set of markers serving as signatures for cell types in *ξ*. Next, a submatrix from matrix *A*, denotes as *A*_<_, containing the intensity values of both the clean cells and the mixed cells, i.e., 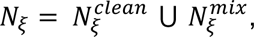 in the marker set *M*_<_ will be extracted. The *k*-nearest neighbors (KNN) algorithm is applied to cell feature matrix *A*_<_ to classify the cells with mixed identities in *ξ*^39^. For each mixed cell in 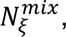 the algorithm works by first calculating its relative distances to clean cells within *ξ*-relevant markers in *M*_<_. This step is crucial as it utilizes only the signature markers for *ξ*, eliminating noise and biases from irrelevant markers in the deconvolution of cell types in *ξ*. The *k* neighbors that are closest to each of the mixed cells will be identified according to their distance. Finally, the identity of a cell is determined by the mode of the identities of its *k*-nearest clean cell neighbors (*k* =10 by default).

### Comparisons with other methods

We compared our proposed method with three existing cell phenotyping methods, namely CELESTA, SCINA, Louvain + manual annotation clusters, and Seurat transfer using scRNA. The code for CELESTA, SCINA, Louvain annotation and Seurat v5 transfers label scRNA methods are publicly available for reproducibility and comparison purposes.

### CELESTA^28^

CELESTA is a cell type identification algorithm for spatial proteomics that uses an optimization framework to assign individual cells to their most likely cell types based on prior knowledge of each cell type’s marker signatures. It utilizes a marker-scoring function to match a cell’s marker expression probability profile to known cell type signatures. In our application, CELESTA was run for each of the tissue microarrays (TMAs). The major function included CreateCelestaObject() to create celesta object. FilterCells() to filter out cells that are artifact, with high_marker_threshold = 0.9, and low_marker_threshold=0.4. AssignCells() function to assigned cell types, with max_iteration=10, and cell_change_threshold=0.01. For each cell type, Additional parameters including high_expression_threshold_anchor, low_expression_threshold_anchor, high_expression_threshold_index, and low_expression_threshold_index need to be defined. As no guidance was provided on how to set the parameters, the default setting was used as provided in this GitHub (https://github.com/plevritis-lab/CELESTA/tree/main). For PCF-HI datasets, CELESTA labeled all cells as Unknown even with the high_expression_threshold_anchor levels were set at 0.2.

### SCINA^13^

SCINA is a method used for cell type identification in scRNA-seq, employing a combination of cell type-specific marker signatures and an expression matrix. Data normalization is performed through log-transformation before further annotation. A signature matrix (referenced in Table S1) is utilized to classify cell types. In the first phase, primary cell types such as vasculature, tumor cells, stroma, immune cells, and smooth muscle are identified. Cells labeled as immune or unknown in the first round undergo a second round of classification, where they are further distinguished into B cells, T cells, CD11c+ dendritic cells, natural killer cells, lymphatics, plasma cells, macrophages, and granulocytes. The third round focuses on cells categorized as T cells or unknown from the second round, aiming to specify subsets like CD4 T cells, CD8 T cells, regulatory T cells (Tregs), and CD45RO+ CD4 T cells. For the PCF-HI, most of the cells return Unknown, therefore, we could not include in the analysis. The SCINA algorithm is executed using the SCINA() function, with parameters such as max_iter = 100, convergence_n = 10, convergence_rate = 0.999, sensitivity_cutoff = 0.9, rm_overlap=TRUE, allow_unknown=TRUE, and log_file=’SCINA.log’. For more information about SCINA, refer to https://github.com/jcao89757/SCINA.

### Louvain^29^

Louvain clustering is a widely used unsupervised method for identifying cell types in spatial omics datasets. This technique, originally developed for community detection in networks, optimizes modularity to partition data into clusters, making it particularly effective for distinguishing distinct cell populations based on gene expression profiles. To run Louvain clustering on spatial omics data, we first normalize the data using z-score normalization to standardize the expression levels. Next, we scale the data to ensure that each feature contributes equally to the analysis. We then perform dimensionality reduction using Uniform Manifold Approximation and Projection (UMAP) on the first 30 principal components to visualize the data in a lower-dimensional space. Finally, we apply Louvain clustering on the UMAP dimensions with a resolution of 0.8 to identify distinct clusters. After that, FindMarkers() function in Seurat version 5 would be used to find the top 5 markers that define the clusters^31^. We look at individual clusters with their expression to assign cell types and the top 5 markers to assign the cell type for each cluster.

### Seurat Label Transfer^31^

Automatic cell labeling was informed by the scRNAseq dataset using post-quality control data. Subsequent data scaling was performed using the ScaleData() function. Dimension reduction was achieved through PCA and UMAP, utilizing the RunPCA() and RunUMAP() functions respectively, focusing on the 30 selected features. The method involved the FindTransferAnchors function from Seurat v5. All 25 clusters remained consistent between the reference (SC) and query (ST) objects.

### Performance metrics

#### Compare with reference

For a specific cell type, True Positive (TP) calls are defined as cells where the assigned cell types from the method match those in the ground truth benchmark dataset. False Positive (FP) calls are cells where the assigned cell types by the method do not match the ground truth or reference. False Negative (FN) calls represent cells assigned by the benchmark but not by the method, while True Negative (TN) calls are cells not assigned by either the method or the benchmark. The weighted score considers the proportion of each cell type in the reference dataset, where *i* is a cell type in the set of reference.

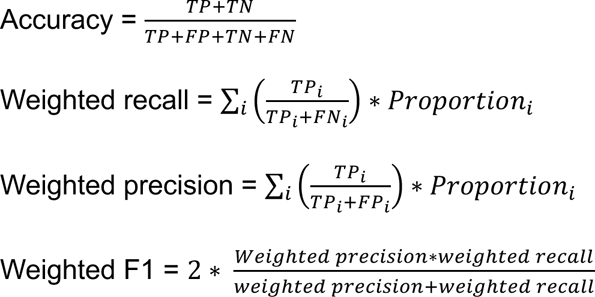

### Benchmark Datasets

Four multiplexed tissue imaging studies with high confidence cell type assignments were used for TACIT evaluation and benchmarking:

<u>PhenoCycler 1 (PCF-Colorectal cancer)</u>^26^: Data representing 140 tissue microarray (TMA) spots from 35 colorectal cancer (CRC) patients (17 in the CLR group and 18 in the DII group) were collected from 36 distinct tissues. In this study, the authors used spatial proteomics to examine the tumor environment and how the immune response correlates with survival outcomes in colorectal cancer. The TMAs were collected and imaged using a 56-marker CODEX (co-detection by indexing) panel, profiling a total of 258,386 cells. Cells identified as immune/vasculature (n=2,153) and immune/tumor (n=1,797), along with cells lacking a marker signature—including adipocytes (n=1,811), nerves (n=659), undefined (n=6,524), monocytes (n=815), and cells categorized as dirt (n=7,357)—were excluded from the analysis. This exclusion resulted in 235,519 cells being retained for the cell type annotation benchmark analysis. The TMA imaging was segmented based on DRAQ5 nuclear stain, pixel intensities were quantified, and spatial fluorescence compensation was performed using the CODEX toolkit segmenter (available at https://github.com/nolanlab/CODEX). Subsequently, the cells were subjected to X-shift clustering, and the resulting clusters were manually annotated to ensure the accuracy of the cell labels. The list of signature was provided in the original paper^26^. PCF-CRC can be download at: https://data.mendeley.com/datasets/mpjzbtfgfr/1.

<u>PhenoCycler 2 (PCF-Human Intestine)</u>^27^: Data from 64 sections of the human intestine were collected from 8 donors (B004, B005, B006, B008, B009, B010, B011, and B012). In this study, the authors used spatial proteomics to examine the structure of the large and small intestines in humans. The raw image data were segmented using either the CODEX Segmenter or the CellVisionSegmenter (available at https://github.com/nolanlab/CellVisionSegmenter). Employing a 57-marker CODEX panel, a total of 2,603,217 cells were profiled. These cells were initially grouped using Leiden clustering and subsequently annotated under the supervision of the authors^40^. To ensure accuracy, the cell type labels were further consolidated by the authors by inspecting back-annotated cell types on the original images. The list of signatures cell types was provided in the original paper and expert domain knowledge. PCF-HI can be download at: https://datadryad.org/stash/dataset/doi:10.5061/dryad.pk0p2ngrf.

<u>MERFISH</u>^30^: The mouse brain datasets include data for 36 mouse sample IDs across a total of 60 slides. In this study, by combining MERFISH with scRNA-seq, we have elucidated the molecular, spatial, and functional organization of neurons within the hypothalamic preoptic region. The raw image data were segmented using a seeded watershed algorithm with DAPI and total mRNA co-stains. Initially, 1,027,848 cells were profiled. These cells were classified using graph-based clustering and subsequently annotated by the authors. For our analyses with TACIT, we excluded 153,080 cells labeled as ‘Ambiguous.’ Additionally, to comply with Louvain’s method requirements, cells where over 70% of genes had zero counts were also removed. The list of signatures cell types was provided in the original paper. After these filtering steps, the dataset prepared for comparison with Louvain includes 505,961 cells covering 170 genes. MERFISH can be downloaded at: https://datadryad.org/stash/dataset/doi:10.5061/dryad.pk0p2ngrf.

<u>Xenium-SjD</u>: A tissue microarray (TMA) was constructed, consisting of 63 cores derived from formalin-fixed paraffin-embedded (FFPE) tissue blocks from 21 patients (11 with Sjögren’s Disease (SjD) and 10 without). Three cores per tissue block were extracted, using a TMA array to organize the blocks, and the patient samples were randomized from 1 to 21. To fit within the fiduciary framework of the TMA, the section was divided in half by scoring, placing 44 cores on a single slide, including 8 additional cores designated for control tissues. The analysis utilized the standard 280-plex Human breast cancer panel according to the protocols provided by 10x Genomics.

<u>Xenium-GVHD</u>: A tissue microarray including three patients with chronic graft-versus-host disease and three healthy minor salivary glands, derived from FFPE tissue blocks, was mounted on a Xenium Slide (10x Genomics). To fit within the fiduciary frame, we melted the original blocks and embedded the samples in one block. The analysis utilized the standard 280-plex human breast cancer panel from 10x Genomics according to the protocol provided by the company.

### Marker Enrichment Strength

For each marker unique to a specific cell type (a marker that is a signature for only one cell type), we calculate the log2 fold change (log2FC) of that marker in the signature cell type compared to the mean value in other cell types where it is not a signature. Additionally, we perform a one-sided Wilcoxon test to determine if the expression of the marker in the signature cell type is significantly greater than its expression in non-signature cell types.

### Statistical Analyses

Statistical analyses were conducted, and figures were created using R (version 4.3.0). For comparisons between two groups, Student’s t-test was used when the assumption of normality was met; otherwise, the non-parametric Wilcoxon rank-sum test was applied. For comparisons involving more than two groups, analysis of variance (ANOVA) was used, followed by post-hoc tests if significant differences were detected. For multiple comparisons, the false discovery rate was used to adjust the P-values (Benjamini-Hochberg procedure). Results were considered statistically significant if P < 0.05 or if the adjusted P < 0.05 for multiple testing.

### Cell-cell interactions and neighborhood analysis

Spatial omics data from each individual tissue was processed that describes cellular interactions as graphs with nodes representing individual cells and edges potential cellular interactions as determined by Delaunay triangulation. A 97^th^ percentile distance threshold was established for each tissue to eliminate edges representing improbably long cell-to-cell distances. Cells classified as "Unknown" (non-deconvoluted cells) were excluded from the analysis before conducting Delaunay triangulation. An interaction matrix was then constructed, with each element *a*_*ij*_representing the number of edges shared between cell type *i* and cell type *j*. To visually represent these differences, a hierarchically clustered heatmap using Euclidean distance was generated.

### Shiny app

The Shiny app (here called **Astrograph**) takes the input of the signature matrix and the CSV file output from TACIT annotation, which includes spatial information, UMAP coordinates, CELLxFEATURE matrix, and marker thresholds. The app provides a user interface with spatial plots and UMAP visuals featuring annotations, marker expression thresholds, and weighted cell type calculations. Users can also access color annotations, spatial neighborhood connections between cell types across the whole tissue or ROI, and Moran’s I for each marker and cell type to identify spatial autocorrelation. Additional tools include annotated mean heatmaps, Voronoi plots, and proportions of cell types and cell state markers.

### Cellenics

The single-cell RNA sequencing dataset was managed, analyzed, and visualized using the Cellenics® community platform (https://scp.biomage.net/) hosted by Biomage (https://biomage.net/). Cellenics® is now Trailmaker, just released by Parse Biosciences. Pre-filtered count matrices were uploaded to Cellenics®. Barcodes were filtered through four sequential steps. Barcodes with fewer than 500 UMIs were removed. Barcodes representing dead or dying cells were excluded by filtering out those with more than 15% mitochondrial reads. A robust linear model was fitted to the relationship between the number of genes with at least one count and the number of UMIs per barcode using the MASS package (v. 7.3-56) to filter outliers. The model predicted the expected number of genes for each barcode, with a tolerance of 1 - alpha, where alpha is 1 divided by the number of droplets in each sample. Droplets outside the prediction interval were removed. The scDblFinder R package v. 1.11.3 was used to calculate the likelihood of droplets containing multiple cells, and barcodes with a doublet score above 0.5 were filtered out. After filtering, each sample contained between 300 and 8000 high-quality barcodes, which were then input into the integration pipeline. Initially, data was log-normalized, and the top 2000 highly variable genes were selected using the variance stabilizing transformation (VST) method. Principal component analysis (PCA) was performed, and the top 40 principal components, explaining 95.65% of the total variance, were used for batch correction with the Harmony R package. Clustering was performed using Seurat’s implementation of the Louvain method. For visualization, a Uniform Manifold Approximation and Projection (UMAP) embedding was calculated using Seurat’s wrapper for the UMAP package. Cluster-specific marker genes were identified by comparing cells of each cluster to all other cells using the presto package’s Wilcoxon rank-sum test. Keratinocytes were isolated from the complete experiment by extracting manually annotated barcodes and filtering the Seurat object. These subset samples were then input into the Biomage-hosted instance of Cellenics®. Filtering steps were skipped since the data was already filtered. The data underwent the same integration pipeline as the full experiment. All cells were manually annotated using relevant literature and CellTypist.

### Ethical Approval

All original research (Figures 3-5; Extended Data 5) complies with country-specific regulations for ethical research engagement with human participants.

Sample Collection and Tissue Preparation: Deidentified minor salivary gland (MSG) tissues were obtained from diagnostic biopsies in healthy and chronic GVHD patients (University of Sao Paulo IRB 65309722.9.0000.0068; MTA 45276721.4.0000.0068 IRB/MTA). All patients seen at the Dentistry Division of the Hospital das Clinicas of Medicine School of University of Sao Paulo reported herein provided informed consent before participation in this research protocol. All patients have received full medical and dental assistance during the research time and will be followed by the oral medicine team unrestricted. Tissues were fixed in a 10% solution of NBF for a minimum of 24h at 4°C and mounted on paraffin-embedded SuperFrost Plus slides (See Supplementary Methods for biopsy and tissue-mounting procedures).

Research participants provided informed consent according to NIH-approved IRB protocols (15-D-0051, NCT00001390) before any study procedures were performed. All participants were assessed and categorized based on the 2016 classification criteria from the American College of Rheumatology (ACR) and the European League Against Rheumatism (EULAR). Comparator tissues were obtained from subjects (non-SjD) who were otherwise healthy and did not meet the 2016 ACR-EULAR criteria. All subjects underwent screening for systemic autoimmunity and received thorough oral, salivary, rheumatological, and ophthalmological evaluations. Clinical investigations adhered to the principles outlined in the Declaration of Helsinki.

#### Clinical Protocol University of Sao Paulo

Patients included in this study were sourced from two distinct pathways. One pathway involved direct inclusion from the São Paulo Capital Death Verification System. This included patients who had died from acute causes and were under 65 years of age. These individuals underwent post-mortem minor salivary gland biopsies within 4 hours of death. Tissue removal was performed using the minimally invasive autopsy technique as described by Matuck et al. (2022) in the Journal of Pathology. The collected tissue samples were then sent to the histology department at the University of São Paulo School of Medicine for further processing as outlined in the described protocol.

GVHD patient biopsies were obtained from the biobank at the University of São Paulo School of Medicine. These patients were re-consented and followed up for chronic GVHD clinical evaluation. The biopsy samples, taken during episodes of oral lesions, were sent to the histology department for processing following the same procedures mentioned above.

#### Spatial Transcriptomics (Xenium) Sample Preparation

The Xenium workflow, using experimental chemistry and prototype instruments and consumables, starts with sectioning 5 μm FFPE tissue sections onto a Xenium slide. These sections are then deparaffinized and permeabilized to make the mRNA accessible. The mRNAs are targeted by the 313 probes and two negative controls: probe controls to assess non-specific binding and genomic DNA (gDNA) controls to confirm that the signal comes from RNA. Probe hybridization takes place overnight at 50 °C with a probe concentration of 10 nM. After a stringency wash to remove un-hybridized probes, the probes are ligated at 37 °C for two hours, during which a rolling circle amplification (RCA) primer also anneals. The circularized probes are then enzymatically amplified (one hour at 4 °C followed by two hours at 37 °C), producing multiple copies of the gene-specific barcode for each RNA binding event, which results in a high signal-to-noise ratio. After washing, background fluorescence is chemically quenched. The biochemistry is designed to minimize autofluorescence, which can be caused by lipofuscins, elastin, collagen, red blood cells, and formalin-fixation. Sections are then placed into an imaging cassette for loading onto the Xenium Analyzer instrument.

#### Spatial Transcriptomics – Xenium

Gene Panel Design: The Xenium in Situ technology employs targeted panels to detect gene expression, this includes 280 genes from the Xenium Human Breast Panel. The probes are designed with two complementary sequences that hybridize to the target RNA and a third region encoding a gene-specific barcode. This allows the paired ends of the probe to bind the target RNA and ligate to form a circular DNA probe. If an off-target binding event occurs, ligation does not happen, which suppresses off-target signals and ensures high specificity.

#### Xenium Analyzer Instrument

The Xenium Analyzer is a fully automated system that includes an imager (with an imageable area of approximately 12 × 24 mm per slide), sample handling, liquid handling, wide-field epifluorescence imaging, capacity for two slides per run, and an on-instrument analysis pipeline. The imager uses a fast area scan camera with a high numerical aperture, a low read noise sensor, and approximately 200 nm per-pixel resolution. Image acquisition on the Xenium Analyzer is performed in cycles. The instrument automatically cycles in fluorescently labeled probes for detecting RNA, incubates, images, and removes them. This process is repeated for 15 rounds of fluorescent probe hybridization, imaging, and probe removal, with Z-stacks taken at a 0.75 μm step size across the entire tissue thickness.

Image Pre-Processing: The Xenium Analyzer captures Z-stacks of images in every cycle and channel, which are then processed and stitched to create a spatial map of the transcripts across the tissue section. Stitching is performed on the DAPI image, taking all stacks from different fields of view (FOVs) and colors to create a complete 3D morphology image (morphology.ome.tif) for each stained region. Lens distortion is corrected based on instrument calibration data, which characterizes the optical system. The Z-stacks are further subsampled to a 3 μm step size, which is empirically determined to be useful for cell segmentation quality. Image features are extracted from overlapping FOVs and feature matching estimates offsets between adjoining FOVs to ensure consistent global alignment across the image. Finally, the 3D DAPI image volumes (Z-stacks) generated across FOVs are stitched together.

#### Multiplex Proteomics (Phenocycler Fusion)

The multiplex analysis was performed on 5 µm FFPE sections mounted on SuperFrost Plus slides (ThermoFisher, MA, USA). The sections underwent deparaffinization and rehydration, followed by immersion in a Coplin jar containing 1:20 AR9 buffer (Akoya Biosciences, MA, USA). The jar was placed in a pressure cooker for 15 minutes at low pressure, then cooled at room temperature for 30 minutes. The samples were then rinsed in deionized water for 30 seconds and in 100% ethanol for 3 minutes. Pre-staining procedures involved immersing the slides in hydration buffer for 2 minutes and staining buffer for 20 minutes (Akoya Biosciences, MA, USA). The primary antibody cocktail was prepared with 4 blockers (G, S, J, and N), each at 9.5 µL in 362 µL of staining buffer. For each slide, 150 µL of the cocktail was aliquoted and 1 µL of each antibody (as listed below) was added. The slides were then placed in a humidity chamber (StainStray, Sigma-Aldrich, MO, USA) and incubated overnight at 4°C. Following incubation, slides were fixed in a post-staining solution for 10 minutes. After fixation, slides underwent sequential 1-minute PBS washes and a 5-minute immersion in ice-cold methanol. The sections were then treated with 200 µL of a final fixative solution for 20 minutes, followed by additional washes to remove the fixative. Slides were dried and mounted using the Akoya flow cell, which seals the flow cell/coverslip onto the slides for 30 seconds. The slides were removed from the press and soaked in 1X PCF buffer (Akoya Biosciences, MA, USA). PCF reporter wells were prepared by covering a 15 mL Falcon tube with aluminum foil, then adding 6.1 mL of nuclease-free water, 675 µL of 10X PCF buffer, 450 µL of PCF assay reagent, and 4.5 µL of concentrated DAPI solution (prepared in-house) to achieve a final DAPI concentration of 1:1000. This reporter stock solution was distributed into 18 amber vials, with each vial containing 235 µL of the solution. For each cycle, 5 µL of reporter was added to each vial, resulting in a total volume of either 245 µL (for 2 reporters) or 250 µL (for 3 reporters) as detailed in Supplemental Methods Table 2. Reporters were selected from Atto550, AlexaFluor 647, and AlexaFluor 750 based on experimental needs. Distinct pipette tips were used to transfer the contents of each amber vial into a 96-well plate. DAPI-containing vials were pipetted into wells in the H-row, while reporter-containing vials were distributed into other rows. Once the wells were filled, they were sealed with adhesive aluminum foil (Akoya Biosciences, MA, USA). Imaging was conducted using a PhenoImager Fusion system connected to a PhenoCycler (PhenoCycler Fusion system from Akoya BioSciences) with a 20X objective lens from Olympus. Solutions required for instrument operation included ACS-grade DMSO from Fisher Chemical, nuclease-free water, and 1X PCF buffer with an added buffer additive. This solution was prepared by mixing 100 mL of 10X PCF buffer, 100 mL of buffer additive, and 800 mL of nuclease-free water.

Antibody List and Reporter List

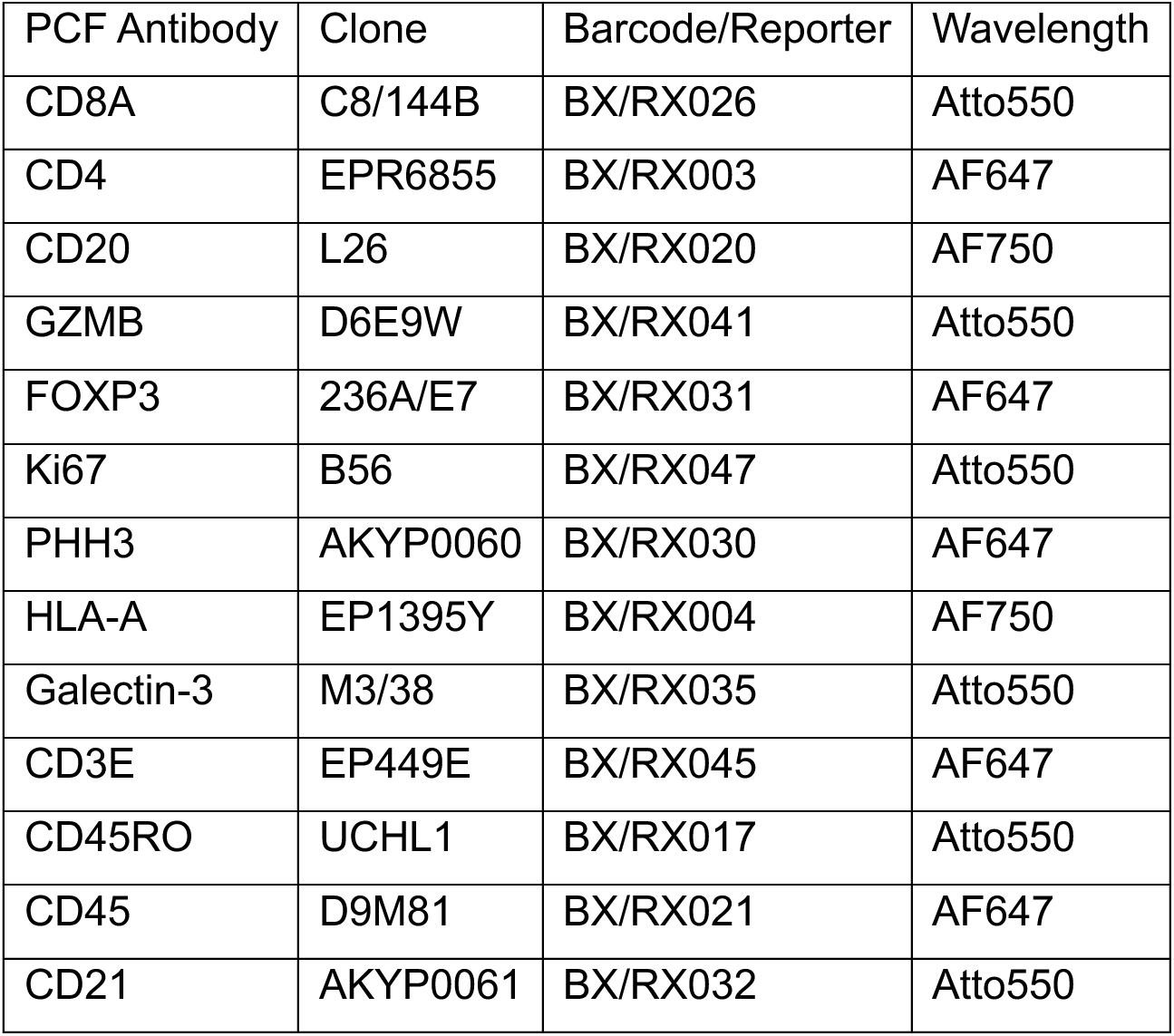
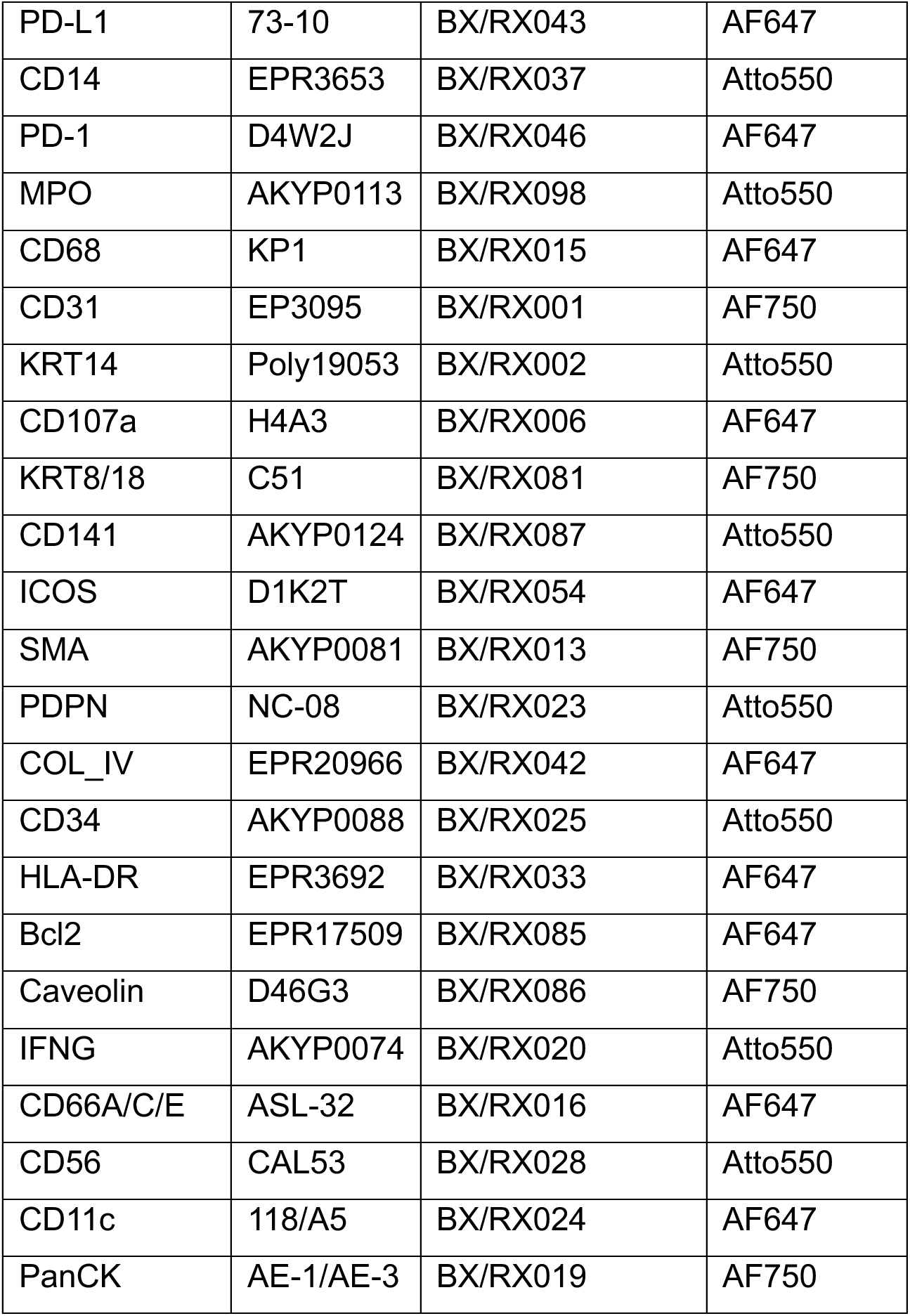

| PCF Antibody | Clone | Barcode/Reporter | Wavelength |
| --- | --- | --- | --- |
| CD8A | C8/144B | BX/RX026 | Atto550 |
| CD4 | EPR6855 | BX/RX003 | AF647 |
| CD20 | L26 | BX/RX020 | AF750 |
| GZMB | D6E9W | BX/RX041 | Atto550 |
| FOXP3 | 236A/E7 | BX/RX031 | AF647 |
| Ki67 | B56 | BX/RX047 | Atto550 |
| PHH3 | AKYP0060 | BX/RX030 | AF647 |
| HLA-A | EP1395Y | BX/RX004 | AF750 |
| Galectin-3 | M3/38 | BX/RX035 | Atto550 |
| CD3E | EP449E | BX/RX045 | AF647 |
| CD45RO | UCHL1 | BX/RX017 | Atto550 |
| CD45 | D9M81 | BX/RX021 | AF647 |
| CD21 | AKYP0061 | BX/RX032 | Atto550 |
| PD-L1 | 73-10 | BX/RX043 | AF647 |
| CD14 | EPR3653 | BX/RX037 | Atto550 |
| PD-1 | D4W2J | BX/RX046 | AF647 |
| MPO | AKYP0113 | BX/RX098 | Atto550 |
| CD68 | KP1 | BX/RX015 | AF647 |
| CD31 | EP3095 | BX/RX001 | AF750 |
| KRT14 | Poly19053 | BX/RX002 | Atto550 |
| CD107a | H4A3 | BX/RX006 | AF647 |
| KRT8/18 | C51 | BX/RX081 | AF750 |
| CD141 | AKYP0124 | BX/RX087 | Atto550 |
| ICOS | D1K2T | BX/RX054 | AF647 |
| SMA | AKYP0081 | BX/RX013 | AF750 |
| PDPN | NC-08 | BX/RX023 | Atto550 |
| COL_IV | EPR20966 | BX/RX042 | AF647 |
| CD34 | AKYP0088 | BX/RX025 | Atto550 |
| HLA-DR | EPR3692 | BX/RX033 | AF647 |
| Bcl2 | EPR17509 | BX/RX085 | AF647 |
| Caveolin | D46G3 | BX/RX086 | AF750 |
| IFNG | AKYP0074 | BX/RX020 | Atto550 |
| CD66A/C/E | ASL-32 | BX/RX016 | AF647 |
| CD56 | CAL53 | BX/RX028 | Atto550 |
| CD11c | 118/A5 | BX/RX024 | AF647 |
| PanCK | AE-1/AE-3 | BX/RX019 | AF750 |

#### Image Segmentation

qpTIFF images were opened into QuPath 5.0, segmentation was acquired in three different methods, the linear nuclei expansion was obtained using Watershed directly from QuPath, the Pre trained models were used applying the QuPath extension generated using the workflow established by Bankhead P. (https://qupath.readthedocs.io/en/latest/docs/advanced/stardist.html).

The HITL methods utilized used a GUI based approach established by Cellpose 3.0 with denoising and HITL training in 50 different ROIs of MSG H&E sections. The application of the methods was performed into a 3 ROIs of 900 microns x 800 microns in three different GVHD patients MSG biopsies. The parameters used by the three methods were the same: Pixel size was 0.1micron, Sigma 1, DAPI threshold 12. Cell expansion 10 into the linear model and the pre-trained model, no cell expansion required for the HITL model. In the HITL model the mask was exported to the QuPath allowing the same extraction csv matrix with the cells IDs and the protein markers expressed in each cell ID.

#### Protocol. Combined Xenium and PCF

After the Xenium experiment, the slides underwent a quenching process as described in the Xenium Assay 10X Genomics manual. The slides were then stored in a container with 50% BPS and 50% glycerol for two days. To resume the experiment, the slide was washed in PBS for 3 minutes, and antigen retrieval was performed using AR9 Buffer (Akoya Biosciences) in a pressure cooker for 15 minutes at low pressure. The rest of the antigen retrieval protocol until the start of the PhenoCycler fusion experiment was carried out as described in the ‘spatial proteomics’ methods section above.

#### Mask Transfer

For the combined Xenium and PCF assay, the cell segmentation masks obtained from Xenium analyzer were used for both Xenium and PCF analysis. Since Xenium acquisition is performed with a 40x objective lens and PCF with a 20x objective lens, for the purposes of cell mask transfer (from Xenium to PCF), the Xenium DAPI image (morpholopgy_mip.ome.tif) was down sampled by a factor of 2. The Xenium cell boundary polygons (stored in cell_boundaries.csv.gz in the Xenium output folder) were subsequently converted to match the downsampled Xenium DAPI image. The cell boundary masks were then saved as a .geojson, with their cell names from Xenium analyzer retained, for use in QuPath for subsequent analysis. Since it is possible for the sample not be perfectly aligned Xenium and PCF experiments, the PCF .qptiff image was registered to the down sampled Xenium DAPI image, using the non-rigid registration workflow in VALIS v1.0.4 (https://www.nature.com/articles/s41467-023-40218-9). The resulting aligned PCF image is saved as an .ome.tiff with the additional downsampled Xenium DAPI channel using the Kheops plugin for FIJI (Guiet, R., Burri, O., Chiaruttini, N., Seitz, A., & Eglinger, J. (2021). Kheops (Version 0.1.8) [Computer software]. https://doi.org/10.5281/zenodo.5256256).

#### Manual Quantification

For the comparison of cell assignment methods, manual counting was conducted by a pathologist (BFM) within designated Regions of Interest (ROIs). These ROIs comprised 1500-1800 cells each. Manual counting involved quantifying cells based on canonical marker labels and morphological features. For example, KRT18 combined with specific morphological features was used to identify Acinar Cells, PAN-Ck combined with morphological features identified Duct cells, CD31 identified Vascular endothelial cells, SMA identified Myoepithelial cells, and CD45 identified immune cells. Additionally, specific markers were utilized for identifying unique cell types that are determined by a single marker. Upon completion of the manual counting process, the quantification data were systematically transferred into a table format. This table facilitated the calculation of the presence of each cell type within the respective ROIs. To assess the convergence between clusters and TACIT, the average number of cells for each type was used to compute the absolute error associated with each cell type.

## DATA AVAILABILITY

The benchmark public data can be found at: https://data.mendeley.com/datasets/mpjzbtfgfr/1 (PCF-CRC), https://datadryad.org/stash/dataset/doi:10.5061/dryad.pk0p2ngrf (PCF-HI), and https://datadryad.org/stash/dataset/doi:10.5061/dryad.pk0p2ngrf (MERFISH). Source data for reproduced figure available at: https://zenodo.org/records/11397609. All other data is available upon reasonable request.

## CODE AVAILABILITY

All codes related to TACIT can be found at https://github.com/huynhkl953/TACIT

## Contributions

For this study, KMB and JL conceptualized the project. JL, KLAH and KMT designed the algorithm. KLAH implemented the algorithm and conducted benchmarking experiments. BFM performed manual assessment of cell type assignment. DDZ, LAVSJ, MD, VGR, LFFDS, DEK, SMH, and BMW supported the recruitment of patients and collected data. KLAH, KMT, BFM, QTE, NVK, TMW, RK, PP, TP, AP, DDZ, KMB, and JL performed experimental and/or bioinformatic analysis that supported project development. JL, KMB, KLAH, KMT and BFM wrote the original draft; KLAH, KMT, BFM, QTE, TMW, DDZ, KMB, and JL critically reviewed and edited the final manuscript.

## Acknowledgements

This work was supported by generous start-up funds from the ADA Science & Research Institute (Volpe Research Scholar Award) and the Chan Zuckerberg Initiative/Foundation program Pediatric Networks for the Human Cell Atlas to KMB. The work also benefited from the VCU Wright Regional Center for Clinical & Translational Science (CCTS) Clinical and Translational Science Award (CTSA) UM1TR004360 to JL and NIH-NCI Cancer Center Support Grant P30 CA016059 to JL and KMT.

## Figure legends

**Extended Data 1:**
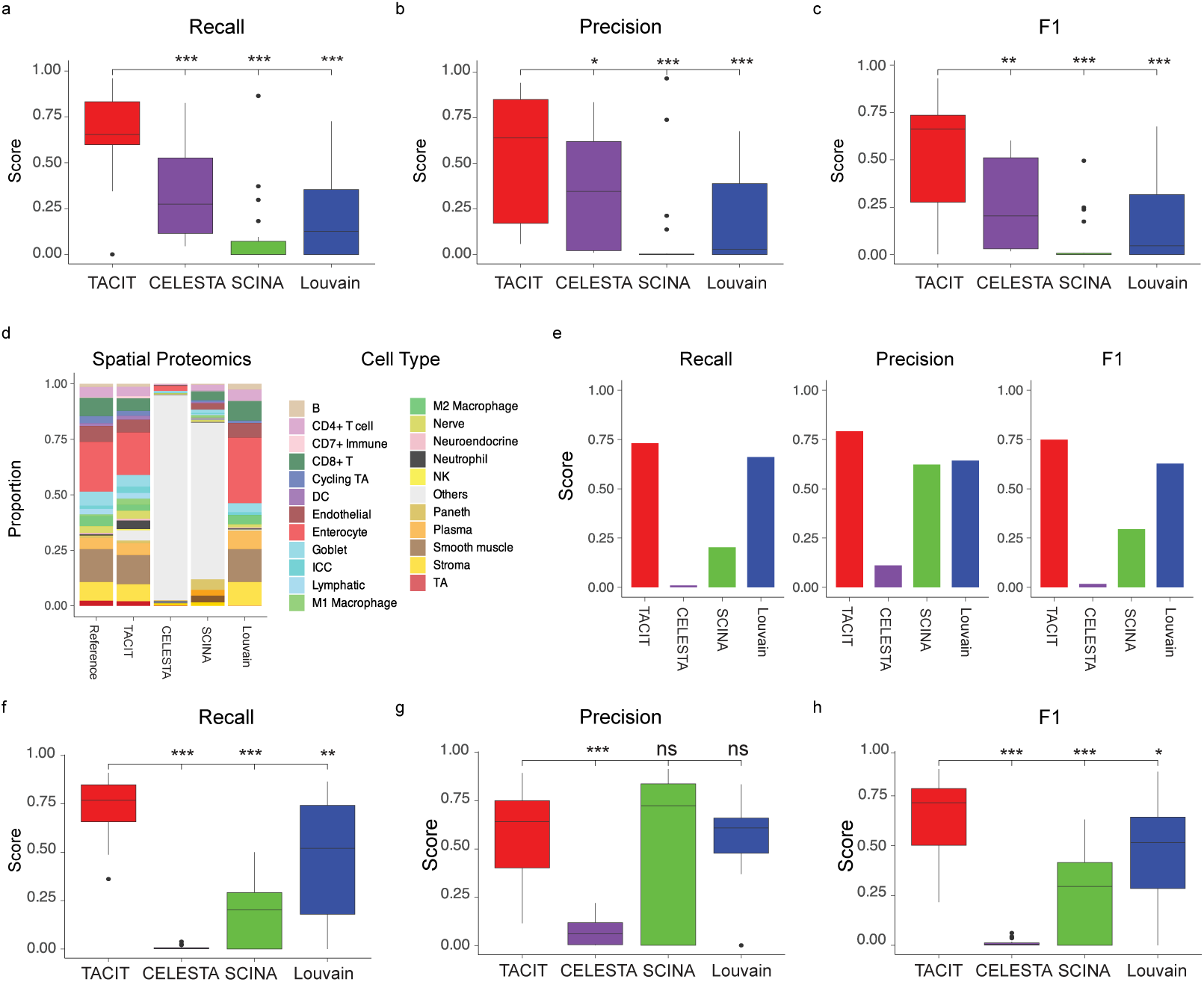
Quantitative of comparison between TACIT and existing methods for individual cell type. (a-c) Boxplots depict recall (a), precision (b), and F1 scores (c) for individual cell types in PCF-CRC, demonstrating the performance of TACIT compared to three alternative methods. TACIT shows significantly higher recall, precision, and F1 scores than CELESTA (p-value<0.05), SCINA (p-value<0.05), and Louvain (p-value<0.05), highlighting its superior accuracy in identifying individual cell types within the PCF-CRC dataset. (d) Comparison of cell type proportions between TACIT and existing methods, with CELESTA and SCINA showing a disproportionately high proportion of the "Others" group in PCF-HI datasets. This over- representation of undefined cell types indicates a limitation in their classification capabilities. (e) Weighted recall, precision, and F1 scores comparing TACIT with existing methods in PCF-HI datasets. Even after excluding the "Others" category, TACIT consistently outperforms other methods, demonstrating higher weighted recall, precision, and F1 scores. (f-h) Boxplots illustrate recall (f), precision (g), and F1 scores (h) for individual cell types in PCF-HI. TACIT’s performance in these metrics remains higher than CELESTA, SCINA and Louvain, further validating its effectiveness in cell type identification.

**Extended Data 2:**
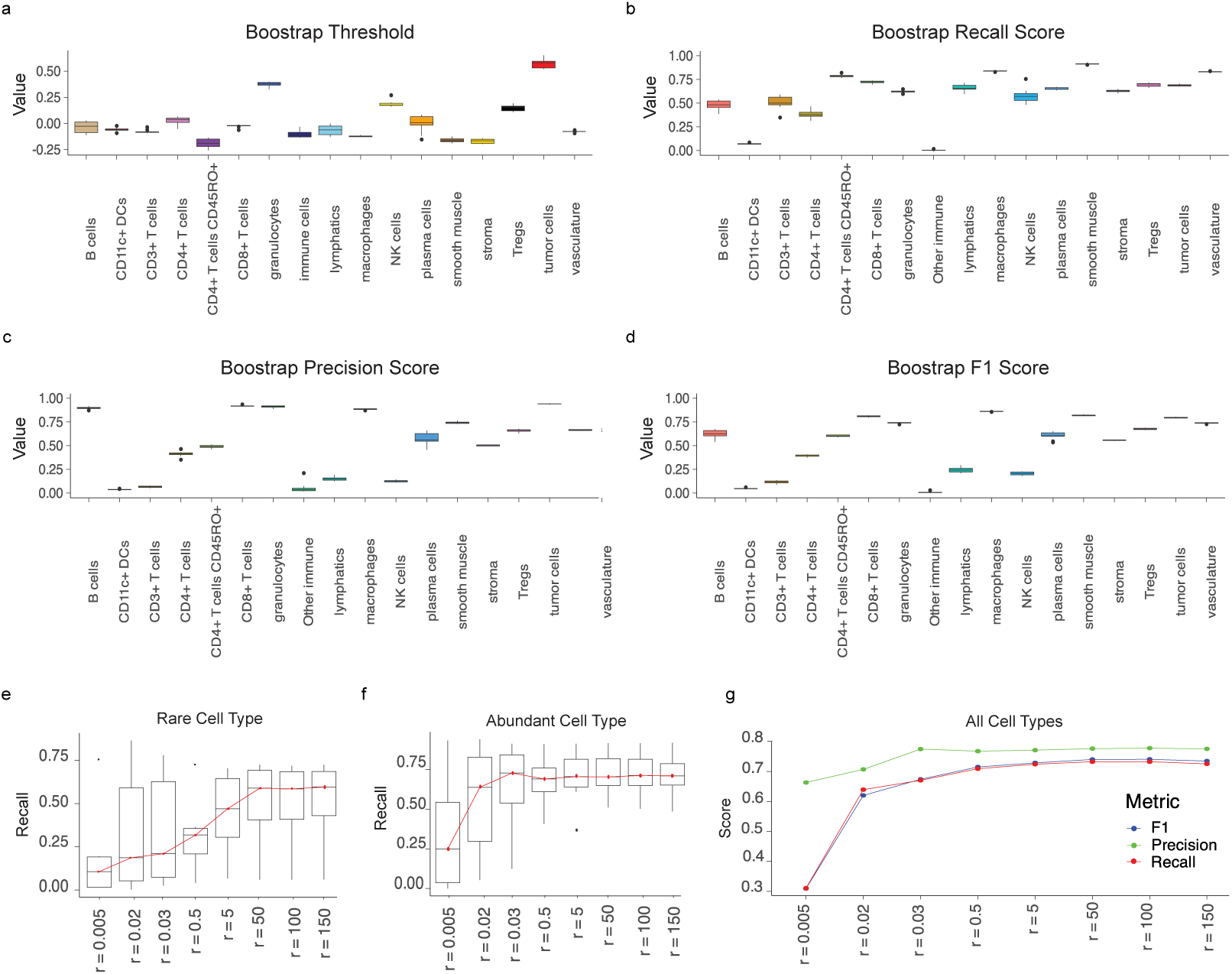
Evaluating Stability and Resolution Impact in TACIT Annotations on PCF-CRC Datasets. Stability and parameter optimization of TACIT annotations are assessed using bootstrap methods on the PCF-CRC datasets. We employed a bootstrap approach by randomly selecting 80% of the original data and running TACIT 10 times to evaluate its stability and robustness. This method ensures that our findings are not biased by any subset of data and provides a comprehensive assessment of TACIT’s performance consistency. (a) The boxplot displays thresholds for each cell type score across the 10 bootstrap iterations, demonstrating that the threshold values remain stable. (b-d) Validation metrics such as recall, precision, and F1 scores are presented for each iteration. These metrics show minimal variation across the 10 bootstrap samples, underscoring TACIT’s reliability in maintaining high performance metrics under different subsets of the data. (e-g) Various resolution levels were tested to assess their impact on the performance of TACIT. Higher resolution levels, which correspond to an increased number of microclusters, showed a positive correlation with recall values, particularly for rare cell types that constitute less than 1% of the data. This enhancement in recall is crucial for accurately identifying and characterizing rare cell populations.

**Extended Data 3:**
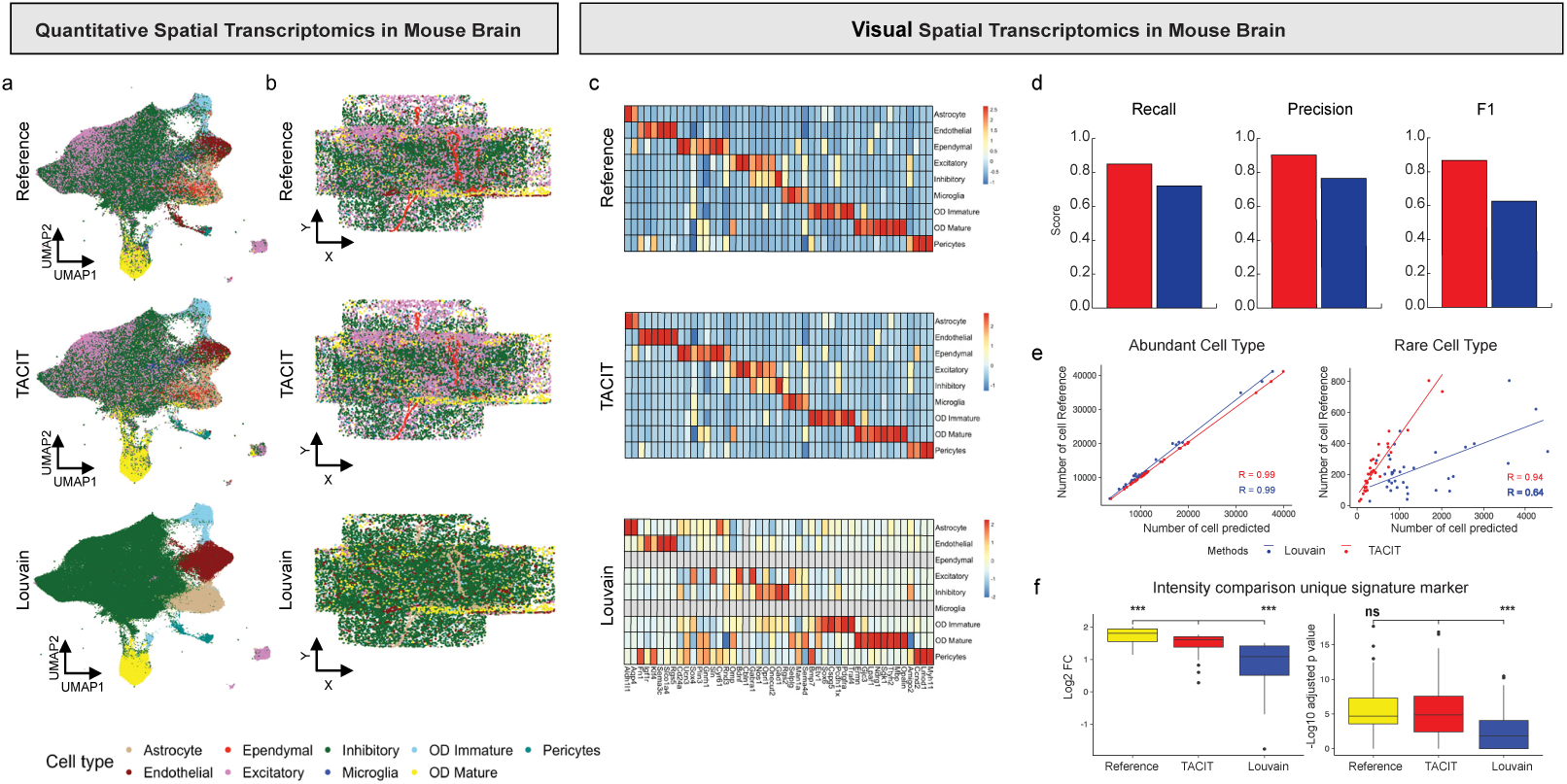
Application of TACIT on MERFISH data from mouse brain. (a) UMAP representations with cell type delineations provide a visual overview of how TACIT effectively clusters cells, showing matching with the reference compared to Louvain clustering. (b) Examples of spatial plots color-coded by identified cell types demonstrate the spatial distribution and organization of cells as identified by TACIT. These plots emphasize TACIT’s ability to preserve spatial integrity, showcasing well-defined structures and consistent cell type placement within the tissue context. (c) Heatmaps comparing the mean marker values for each cell type identified by TACIT and Louvain, along with provided reference data, illustrate the distinct marker expression patterns for each cell type. (d) Comparison of weighted recall, precision, and F1 scores between TACIT and Louvain, benchmarked against the reference, demonstrates TACIT’s superior performance. TACIT consistently achieves higher scores across these metrics (Recall = 0.85, Precision = 0.87, and F1 = 0.85). (e) Correlation plots illustrating the relationships between different cell type identification methods for both abundant (R=0.99) and rare cell types (R=94) reveal TACIT’s strong correlation with reference data. (f) Intensity comparison of unique markers between TACIT and existing methods shows that TACIT exhibits higher intensities of unique marker expressions, which log2FC and -log10 adjusted p-value significant different than Louvain (p-value<0.05).

**Extended Data 4:**
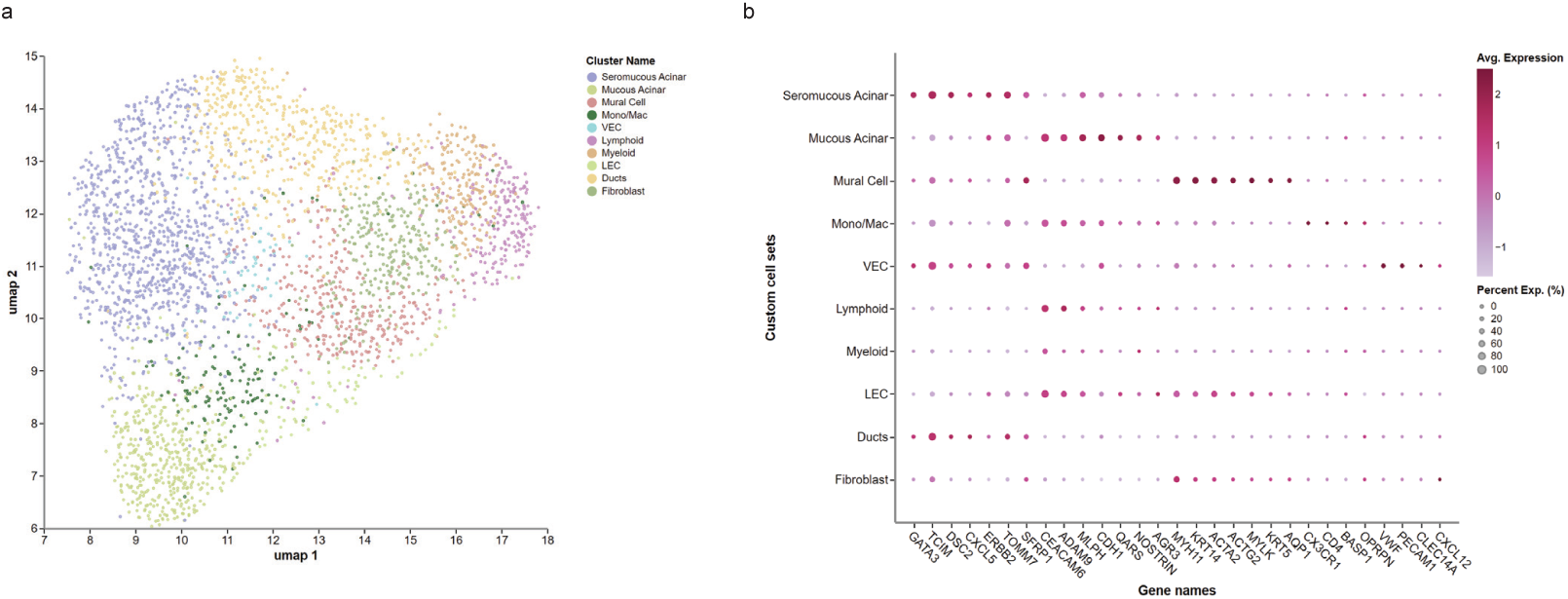
Using biased cell annotation to discover cell types signatures to support spatial analyses. (a) Trailmaker (Parse Biosciences; formerly, Cellenics®) was used to perform cell type annotation on UMAPs. (b) Differentially expressed genes that become a marker set per cluster were generated using their dot plot tool.

**Extended Data 5:**
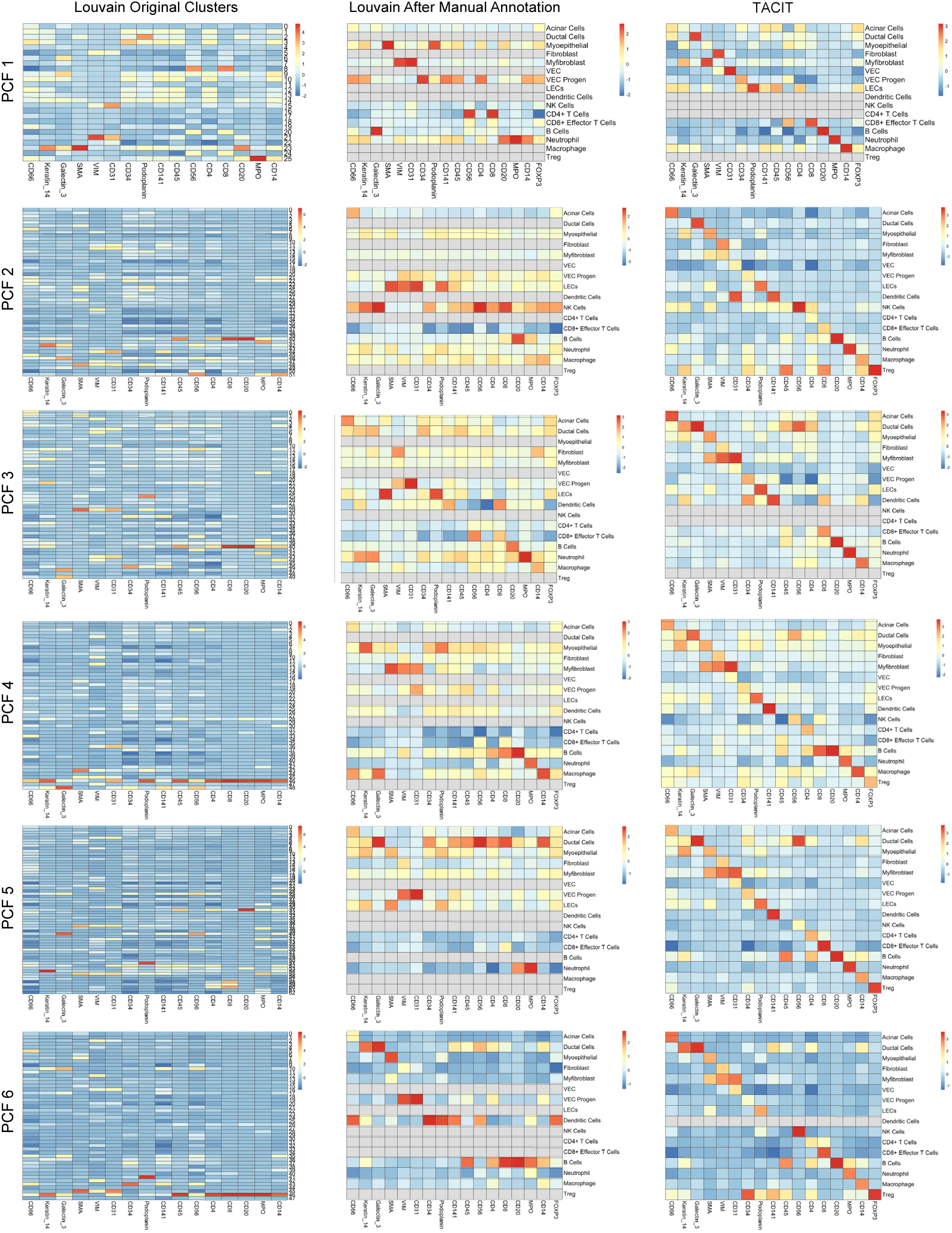
Heatmap of PCFs in 6 Xenium and PCF paired samples. The heatmap displays the mean expression of each antibody and cell type, annotated using three methods: Louvain with default resolution = 0.8 (first column), Louvain after manual annotation (second column), and TACIT annotation (third column). For the first column, we used Louvain clustering with a default resolution of 0.8. In the second column, we manually annotated the Louvain clusters by examining the different gene expressions and identifying the top three markers for each cluster. We then compared these markers with known signatures to assign cell types to each cluster. In the third column, we present the results from TACIT annotation. When comparing TACIT to the Louvain-based methods, TACIT provides more unique and clear diagnostic markers, resulting in distinct and well-defined cell type annotations. This clarity and uniqueness in marker expression underscore TACIT’s superior performance in accurately identifying cell types.

**Extended Data 6:**
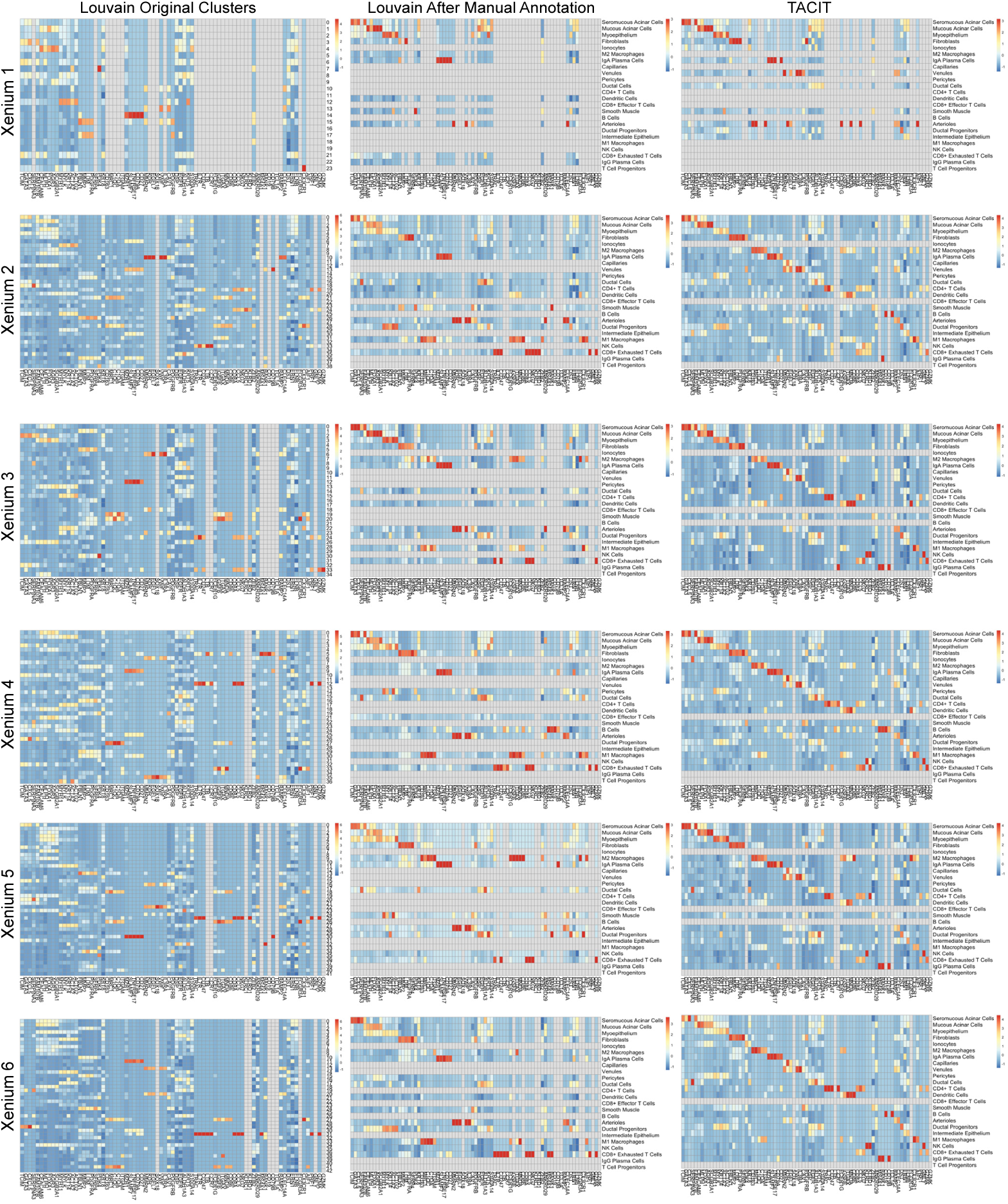
Heatmap of Xeniums in 6 Xenium and PCF paired samples. The heatmap displays the mean expression of each antibody and cell type in the Xenium dataset, annotated using three different methods: Louvain with default resolution = 0.8 (first column), Louvain after manual annotation (second column), and TACIT annotation (third column). For the first column, we utilized Louvain clustering with a default resolution of 0.8. In the second column, the Louvain clusters were manually annotated by examining the different gene expressions and identifying the top three markers for each cluster. These markers were then compared with known signatures to assign specific cell types to each cluster. In the third column, we show the results from TACIT annotation. Compared to the Louvain-based methods, TACIT delivers more distinct and unique diagnostic markers, leading to clearer and more precise cell type annotations. This enhanced clarity and uniqueness in marker expression highlight TACIT’s superior capability in accurately identifying cell types within the Xenium dataset.

**Extended Data 7:**
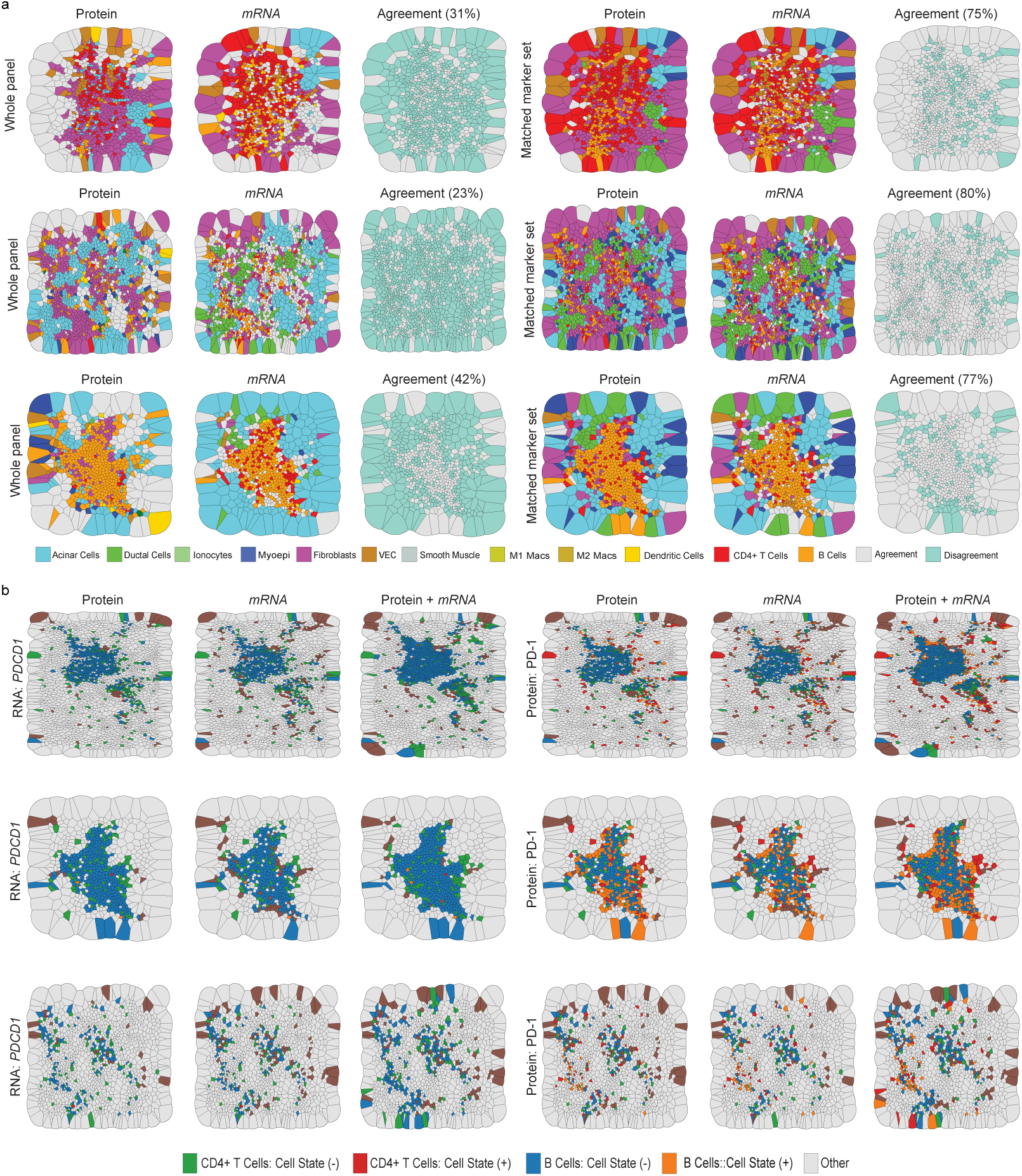
Multimodal analysis between spatial transcriptomics and spatial proteomics in TLS region. (a) TLS region with initial TACIT annotation using the whole panel of Xenium and PCF versus TACIT annotation using matched marker sets. When using the entire panel, the agreement between the two technologies (Xenium and PCF) was relatively low, with agreement rates of 31%, 23%, and 42%. However, focusing on matched marker sets significantly improved the agreement to 75%, 80%, and 77%, respectively, between PCF and Xenium. (b) The cell state (*PDCD1* and PD-1) expression with cell type in the TLS region. This highlights the differences in *mRNA* and RNA cell states, providing insights into the expression patterns and potential discrepancies in cell type identification based on different technologies.

**Extended Data 8: Signature matrix for all datasets.** The input signature matrix for all datasets used in this manuscript.

